# A large but weakened soil carbon sink in resampled forests and grasslands

**DOI:** 10.1101/2025.04.25.650716

**Authors:** Ruofei Jia, Evan C. Fricke, Avni Malhotra, Yinon M. Bar-On, Jie Deng, Gervasio Piñeiro, Bruno Bazzoni, Roberto Alvarez, Nicola Findlay, Mariska te Beest, Yong Zhou, Thomas W. Boutton, Javid Ahmad Dar, Subashree Kothandaraman, Andrew S. MacDougall, Nico Eisenhauer, Elizabeth T. Borer, Eric W. Seabloom, Pablo L. Peri, Jianqiu Zheng, Sally A. Power, Sasha C. Reed, Petr Macek, Sylvia Haider, Hector Bahamonde, Anu Eskelinen, Stephen Sitch, Michael O’Sullivan, Pierre Friedlingstein, Ben Bond-Lamberty, Bruce A. Hungate, Robert B. Jackson, Mina Subramanian, Kaizad Patel, César Terrer

## Abstract

The land carbon sink is composed of vegetation and soil carbon change, but only vegetation change has been quantified by observations at global scale (*1–6*). Recent changes in soil organic carbon (SOC) remain highly uncertain, with process models disagreeing even on its direction (*7*, *8*). Repeated soil sampling is the most direct measure of SOC change, but these measurements have not been synthesized. Here, we compiled repeated SOC measurements from 4,883 sites, representing environmental conditions across 30% of the world’s forests and grasslands without land-use change. Within this domain, soils gained 0.75 [95% credible interval: 0.50, 1.03] petagrams of carbon per year (Pg C yr^-1^) from 1992 to 2020, primarily driven by boreal old forests, temperate young forests, and temperate grasslands. This SOC sink was the highest in the 2000s (1.06 [0.65, 1.48] Pg C yr^-1^), but weakened by 61% in the 2010s (0.41 [-0.13, 0.97] Pg C yr^-1^), as boreal old forests shifted from a sink to a source. The SOC sink is roughly seven times the mean of 17 process models and corresponds to 83±32% of the modeled land sink over the same area. These results suggest that soils account for most of the modeled land sink in the area represented by repeated soil measurements, and they provide an observational benchmark for SOC change in process models.

---

The terrestrial biosphere currently sequesters approximately 20% of anthropogenic carbon dioxide (CO_2_) emissions, serving as a critical buffer against climate change (*9*). This land carbon (C) sink is driven by global change factors excluding the direct effects of land-use change. Observations and models indicate the land sink has increased since the 1990s, largely because the C gains from CO_2_ fertilization and nitrogen (N) deposition have, on average, outpaced climate-driven C losses (*10*, *11*). Yet, whether the land sink will persist or become a source remains uncertain (*12*). A key reason is that models used to project the future land sink do not agree on how the current sink is partitioned between vegetation and soils (*13*). Because carbon resides longer in soils than in vegetation (*14–16*), a model can reproduce the current land sink with an incorrect partition between the two pools and still project the future sink incorrectly. Reliable land sink projections therefore require knowledge of the recent changes in both vegetation and soil carbon.

While changes in vegetation carbon have been quantified from observations at the global scale (*1–6*), changes in soil organic carbon (SOC) have not (*17*). The two existing estimates of global-scale SOC change both use indirect methods and disagree on the soils’ contribution to the land sink. A forest inventory study attributed about 10% of the forest C sink to soils, but it estimated SOC change primarily with models because soils lack remeasurements (*2*). A second analysis inferred that non-living pools (soil, litter and woody debris) stored more than half of the recent C gain on land (*5*). This estimate is also not derived from SOC measurements, but the difference between the total C gain on land and the other terms of the budget (living vegetation, sediment transport, harvested wood products and landfills). It therefore inherits the uncertainties of all these terms and does not separate SOC from litter and woody debris. For example, satellites may underestimate C accumulation in mature forests (*18*), in which case the gain in non-living pools is overestimated.

The direction of recent SOC change remains unknown. CO_2_ fertilization of plant productivity may have increased SOC (*10*, *19*), and warming may have decreased it through higher heterotrophic respiration (*20–22*). Process models agree on the sign of each effect (*23*) but disagree on the direction, magnitude and spatial pattern of global SOC change (*7*, *8*). It is also unknown whether SOC change has been stable over recent decades. Evidence from the vegetation sink suggests such instability is plausible: the C sink of intact Amazonian forests has declined since the 1990s, whereas the sink of intact African forests remained stable (*4*). These patterns in vegetation change were detected by measuring the same forest plots repeatedly (*3*, *4*). For soils, repeated sampling is also the most direct measure of SOC change, but these measurements have not been synthesized.

In this study, we assembled repeated SOC measurements from 4,883 forest and grassland sites without land-use or land-cover change. Each site was sampled at least twice between 1957 and 2022 (Fig. 1). First, we fitted a Bayesian generalized additive mixed model (GAMM) (*24*) to estimate SOC trends in the top 30 cm. Then, we applied it to the natural systems within the environmental conditions represented by the observations. Here, we define “natural systems” as uncultivated ecosystems that did not experience land-use change during the studied period, such as forests remaining forests and grasslands remaining grasslands. Finally, we compared the resulting observational SOC sink with the modeled SOC sink and land sink.

**Fig. 1.**
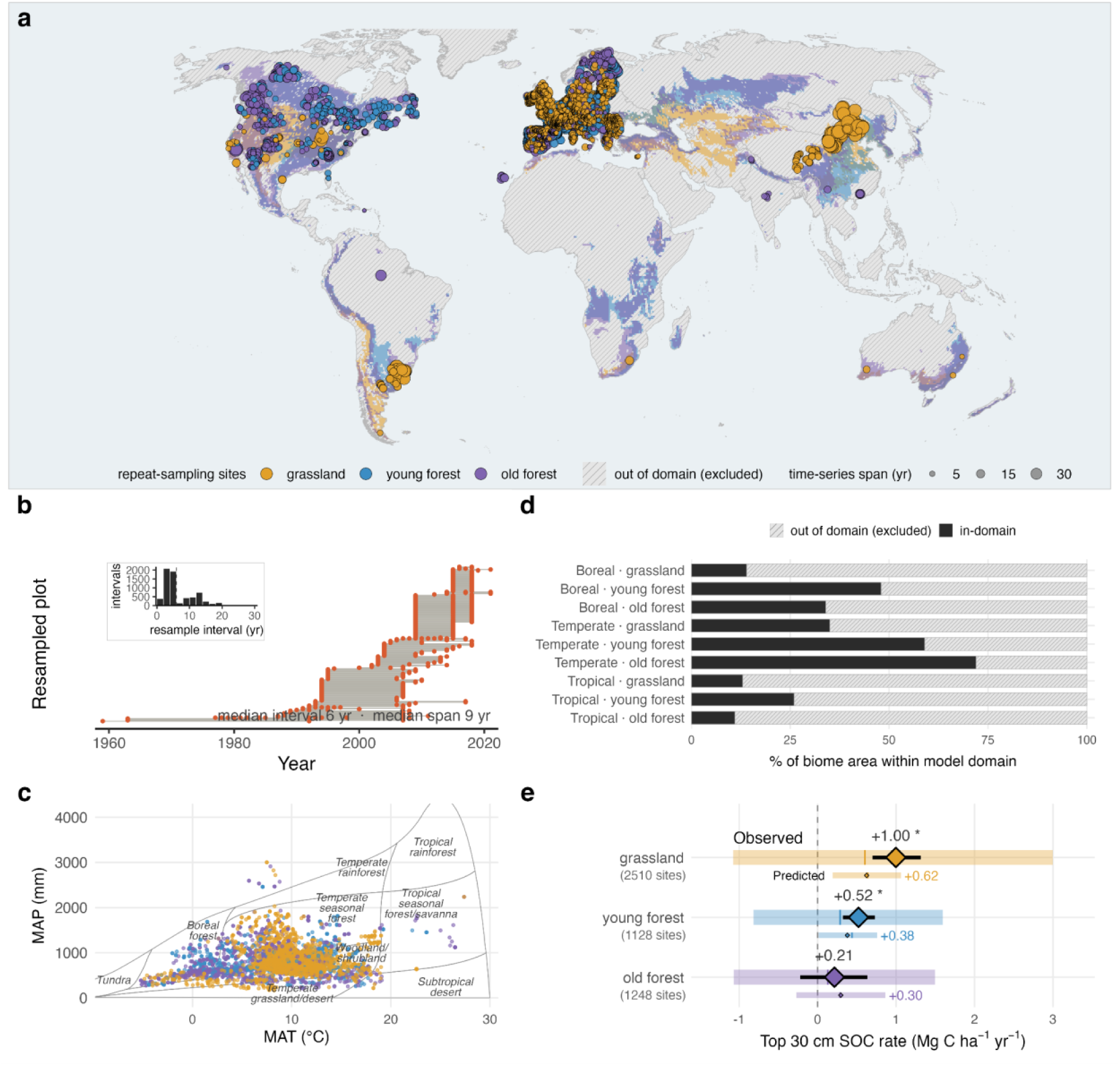
Database of repeated soil measurements and the spatial domain of analysis. (a) Spatial distribution of the database. The points represent 5,364 SOC time series, colored by land-cover class and sized by the time span of each time series. Shaded areas indicate the spatial domain of upscaling, where all pixels lie within the environmental conditions of input predictors. Hatching indicates out-of-domain areas that are not evaluated by this analysis. (b) Temporal structure of the database visualized by 180 random time series. Each row is one plot-level time series and points are individual visits. The inset shows the distribution of resample intervals. (c) Climate space of the sampled plots over Whittaker biome boundaries. (d) Percentage of each biome’s applicable area that falls within the analysis domain. The climate zone division follows Extended Data Table 1. (e) Descriptive statistics of observed and predicted pixel-level SOC rates. Both evaluate SOC change at the top 30 cm. Shaded bars and vertical lines indicate the 25%–75% percentile range and median values. For observed SOC rates, diamonds and whiskers indicate the mean values and their 95% bootstrap confidence intervals (CI). Annotated mean values have asterisks if the 95% bootstrap CI does not overlap with zero. Statistics for observed SOC rates were weighted by the inverse of replicate plot number at each site. The site numbers from each land cover do not add up to the total 4,883 because three sites have both forest and grassland plots. For predicted SOC rates, diamonds and annotated values indicate the average of posterior mean predictions of decadal SOC rates on all in-domain pixels, during 1992–2000, 2000–2010, and 2010–2020. Whiskers and asterisks are not applicable to the predicted SOC rates.

## A database of repeated soil sampling

The database contains 5,364 plot-level SOC time series from 4,883 forest and grassland sites sampled between 1957 and 2022, resulting in 12,319 samplings in total (Fig. 1a). All sites experienced no land-use or land-cover change during the sampling time span. Plots were revisited at a median interval of 6 years, and the median time series span is 9 years (Fig. 1b). A quarter (1,339) of the time series were sampled three or more times, up to nineteen. Sampled sites span mean annual temperatures from -5.2 to 27.4 °C and precipitation from 167 to 3,003 mm (Fig. 1c), and are mostly located in Europe, North America, China, and Pampas grasslands in South America.

We restricted the estimates of recent SOC change to the area with climate and soil conditions represented by the observations (Figs. 1a, 1d, Supplementary Figs. S1 and S2). The model is therefore not extrapolated beyond the sampled conditions. Pixels (0.5°×0.5°) were included if their climate and soil conditions lie within the range of the sampled sites without outliers (in-domain; colored in Fig. 1a). All other pixels were excluded (out-of-domain; hatched in Fig. 1a). This analysis domain covers 30% of the global area of forests and grasslands without land-use change. Depending on land covers, this domain spans 14–48% of boreal biomes, 35–72% of temperate biomes, and 11–26% of tropical biomes (Fig. 1d, Extended Data Table 1). The in-domain tropical areas comprise higher-latitude and montane regions of South America and Africa, and exclude tropical moist forest ecosystems. In-domain grasslands are mostly limited to the temperate regions, with boreal and tropical grasslands taking up only <5% of the total in-domain area (Supplementary Fig. S1). Some regions with few or no local sites are within the domain because their climate and soil conditions are represented by sites in other regions. These include Southern Russia, Central Asia, Oceania, and higher-latitude and montane South America and Africa. Note that results on these regions are interpolated from fitted trends and may not account for local conditions beyond those captured by predictors.

Across 4,883 forest and grassland sites, SOC stocks increased at an average rate of 0.69 [95% confidence interval: 0.48, 0.89] Mg C ha^-1^ yr^-1^ in the top 30 cm. By land cover (Fig. 1e), SOC increased by an average of 1.00 [0.70, 1.31] Mg C ha^-1^ yr^-1^ in grasslands and 0.52 [0.32, 0.73] Mg C ha^-1^ yr^-1^ in young forests (<60 years). In old forests (>=60 years), the average SOC change was 0.21 [-0.22, 0.63] Mg C ha^-1^ yr^-1^), which does not differ from zero. These mean rates, however, only describe the SOC changes on sampled locations (Fig. 1a). Statistical modeling is required to estimate SOC changes across all forests and grasslands in the study domain and to propagate their uncertainty to broad-scale averages and totals.

## Recent SOC sink in forests and grasslands

To estimate SOC change over area and time, we fitted a Bayesian GAMM to all visits of the 5,364 time series. The model predicts the top 30 cm SOC stock from different years based on climate, soil properties and land cover, so the estimate for each pixel depends on its environmental conditions rather than location. The model allows for nonlinear temporal trends, so changes in SOC sink between decades can be estimated directly. Site and plot random intercepts account for spatial variation in SOC across time series, allowing temporal trends to be informed by within-site change rather than site-to-site difference. The model’s posterior distribution quantifies the uncertainty of SOC sink or source estimates at each scale, from individual pixels to the whole domain, propagated consistently from the same posterior samples. The converged model explained substantial variation in the SOC stock across observations (marginal R^2^=0.49, Methods, Supplementary Fig. S3, Table S1) (*25*). The model’s posterior mean predictions of SOC rates on in-domain pixels fall within the observed ranges and cluster near the observed medians (Fig. 1e).

Within the analysis domain, forest and grassland soils gained 0.75 [95% credible interval: 0.50, 1.03] Pg C yr^-1^ during 1992–2020, with more than 99% posterior probability of a sink (Fig. 2, Extended Data Figs. 1–5, Extended Data Table 2). The accumulative gain was 21.08 [14.10, 28.83] Pg C. This SOC sink weakened in the most recent decade (Fig. 2, Extended Data Figs. 6, 7). It was 0.79 [0.33, 1.25] Pg C yr^-1^ in the 1990s and 1.06 [0.65, 1.48] Pg C yr^-1^ in the 2000s, with no detectable change between them (79% posterior probability of an increase). In the 2010s, it was 0.41 [-0.13, 0.97] Pg C yr^-1^, 61% lower than in the 2000s (96% posterior probability of a decrease). Boreal old forests account for almost all of this decrease. They changed from a SOC sink of 0.33 [0.07, 0.59] Pg C yr^-1^ in the 2000s to a source of -0.30 [-0.59, -0.02] Pg C yr^-1^ in the 2010s. This difference of 0.63 Pg C yr^-1^ is 97% of the total decrease of 0.65 Pg C yr^-1^. The SOC sink in young forests did not change detectably between decades, and the sink in grasslands increased from the 1990s to the 2000s (99% posterior probability).

**Fig. 2.**
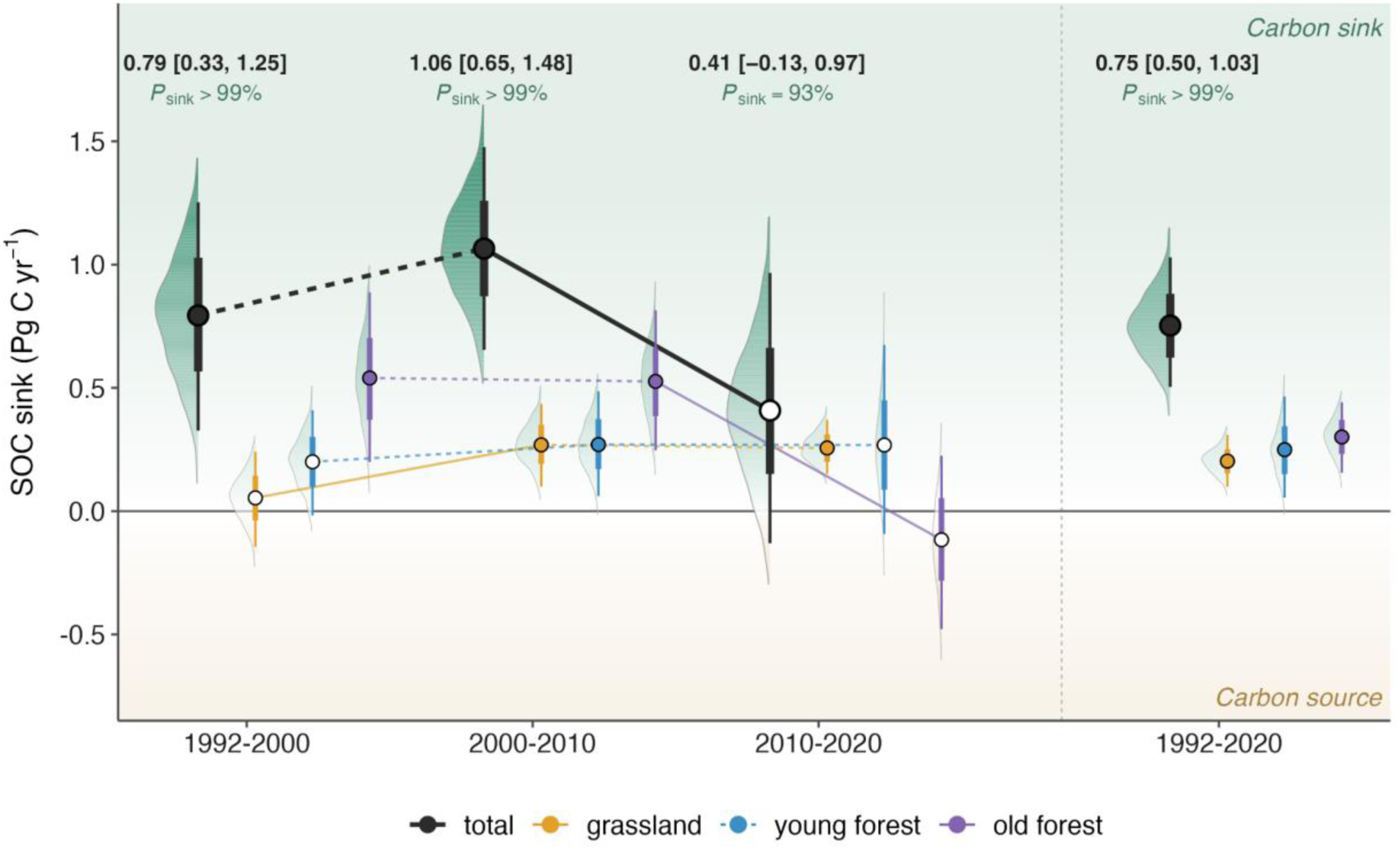
Magnitude and recent weakening of the natural SOC sink. Half violins indicate posterior predictive distributions of the annual SOC sink (Pg C yr^-1^) in each recent decade (1992–2000, 2000–2010, 2010–2020) and over all three decades (1992–2020; right of the divider). Distribution tails were trimmed where posterior density falls below 3% of the peak. Circles indicate posterior means, and the thick and thin bars indicate the 66% and 95% credible intervals. Annotated sink values are mean and [2.5%, 97.5%] percentile of the posterior estimates. *P*_sink_ above each estimate indicates the posterior probability that the SOC change is positive. Filled circles indicate *P*_sink_ ≥ 97.5% and hollow circles indicate *P*_sink_ < 97.5%. Solid lines between two decadal estimates indicate that the SOC sink has strengthened or weakened with more than 95% posterior probability. Dashed lines indicate no detectable strengthening or weakening of the SOC sink (<95% posterior probability).

All three land covers contributed to the natural SOC sink during 1992–2020: old forests contributed 40%, young forest 33%, grasslands 27%. These percentages roughly mirror each land cover’s share of the analysis domain (48%, 31%, 20%, respectively). The three strongest biome-level sinks were boreal old forests (0.2 [0.09, 0.31] Pg C yr^-1^), temperate grasslands (0.16 [0.08, 0.24] Pg C yr^-1^), and temperate young forest (0.12 [0.05, 0.19] Pg C yr^-1^). Together, they account for 64% of the total sink (Extended Data Table 2). Although temperate old forests cover 13% of the total in-domain area, they contributed only 6% (0.05 [-0.004, 0.09] Pg C yr^-1^) of the SOC sink, because their SOC loss in the 1990s offset accumulations in later decades (Fig. 3, Extended Data Table 2). Regionally, boreal old forests in Canada and Russia have the largest SOC gains over 1992–2020 because of their high accumulation rates and large area, which covers 22% of the analysis domain (Fig. 3, Extended Data Tables 1–3). In both regions, the sinks weakened from the 2000s to the 2010s with more than 95% posterior probability, and became SOC sources in the 2010s (Extended Data Table 2). Young forest SOC sinks are the largest in East Asia because of its vast young forest area included in the analysis. Grassland SOC sinks are concentrated in the United States, Europe, Central and West Asia, and East Asia, in that order, reflecting both the large in-domain grassland area and high accumulation rates in these regions (Extended Data Tables 2, 3).

**Fig. 3.**
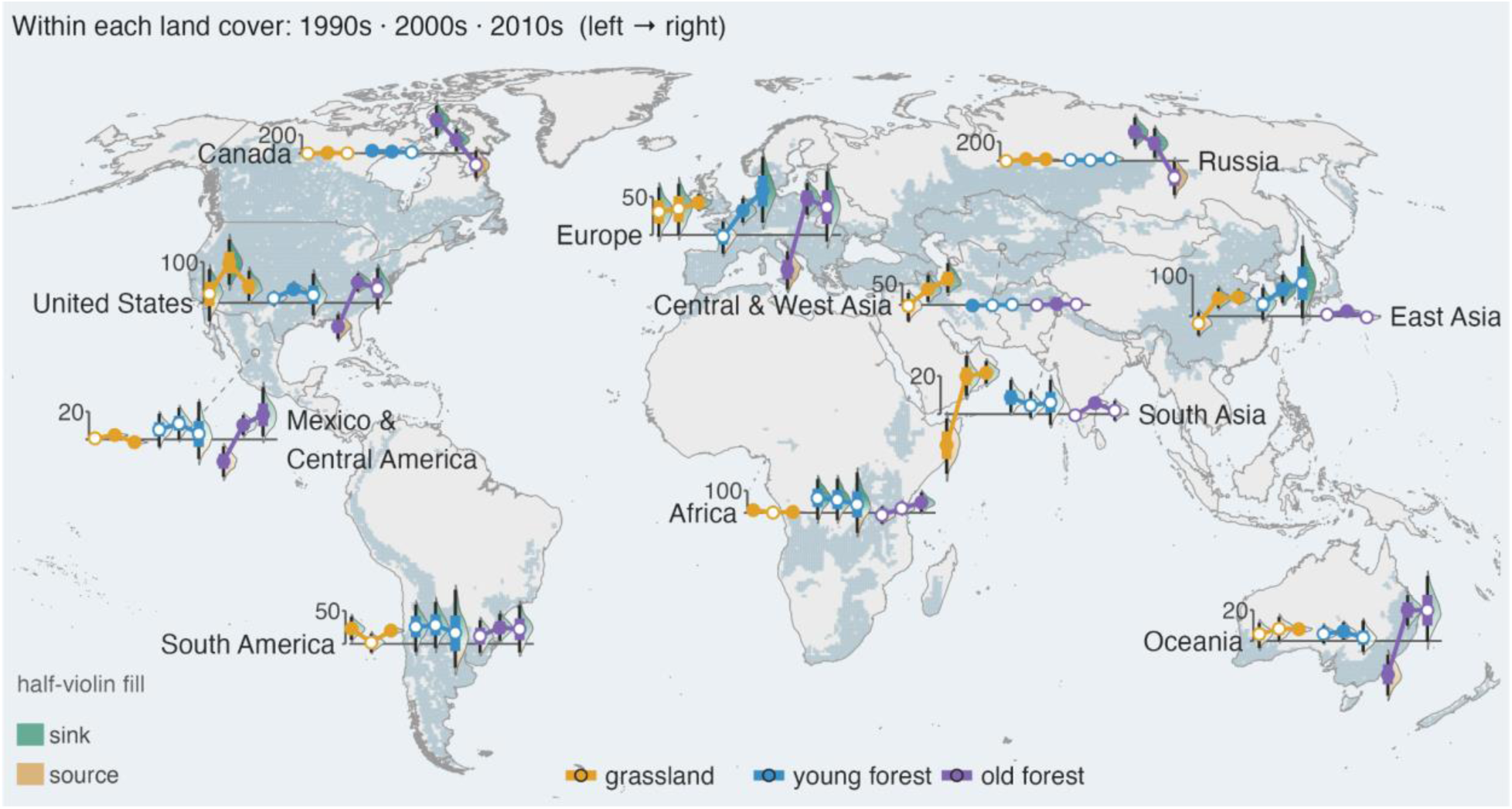
Regional estimates of the natural SOC sink (Tg C yr^-1^). Half violins indicate the posterior predictive distributions of decadal SOC sinks (1992–2000, 2000–2010, 2010–2020). Circles indicate posterior means, and the thick and thin bars indicate the 66% and 95% credible intervals. Distribution tails were trimmed where posterior density falls below 6% of the peak. Filled circles indicate a SOC sink with more than 97.5% posterior probability. Estimates of Alaska, Southeast Asia, and Northern Europe are omitted in this figure but reported in Extended Data Table 2. Regional SOC sinks over all three decades (1992–2020) and the strengthening or weakening of regional sinks are reported in Extended Data Table 2.

## Comparisons with modeled SOC sink and land sink

This analysis provides an observational benchmark for the divergent model estimates of the recent SOC sink (*7*, *8*), as well as its contribution to the land sink (*13*). Here, we compared the in-domain SOC sink in old forests and grasslands during 1992–2020 with the modeled SOC sink from TRENDY-v12 S2 simulations (*26*) and the modeled land sink from the Global Carbon Budget (GCB) (*9*). We excluded young forests from the comparisons because the modeled SOC sink and land sink both classify C gains in young forests as part of land-use change emissions rather than the land sink. Without young forests, the in-domain SOC sink is 0.50 [0.32, 0.68] Pg C yr^-1^. We used this value to compare with the models.

Over the same spatial and temporal domain of this analysis, models from TRENDY-v12 simulate a SOC sink of 0.07 ± 0.06 Pg C yr^-1^ (mean ± SD across 17 models; range [-0.04 to 0.18], Supplementary Table S2). Our observation-based estimate (0.50 [0.32, 0.68] Pg C yr^-1^) is roughly seven times the ensemble mean, and every model is below the lower bound of its 95% credible interval (Fig. 4a). The discrepancy is likely understated, because the models integrate the full soil profile while the observation-based estimate covers only the top 30 cm. This mismatch suggests that the SOC sink may be higher than represented by process models across the analysis domain. The database of repeated soil sampling (Fig. 1) provides an opportunity for optimizing process models with direct SOC change measurements (*27*, *28*).

**Fig. 4.**
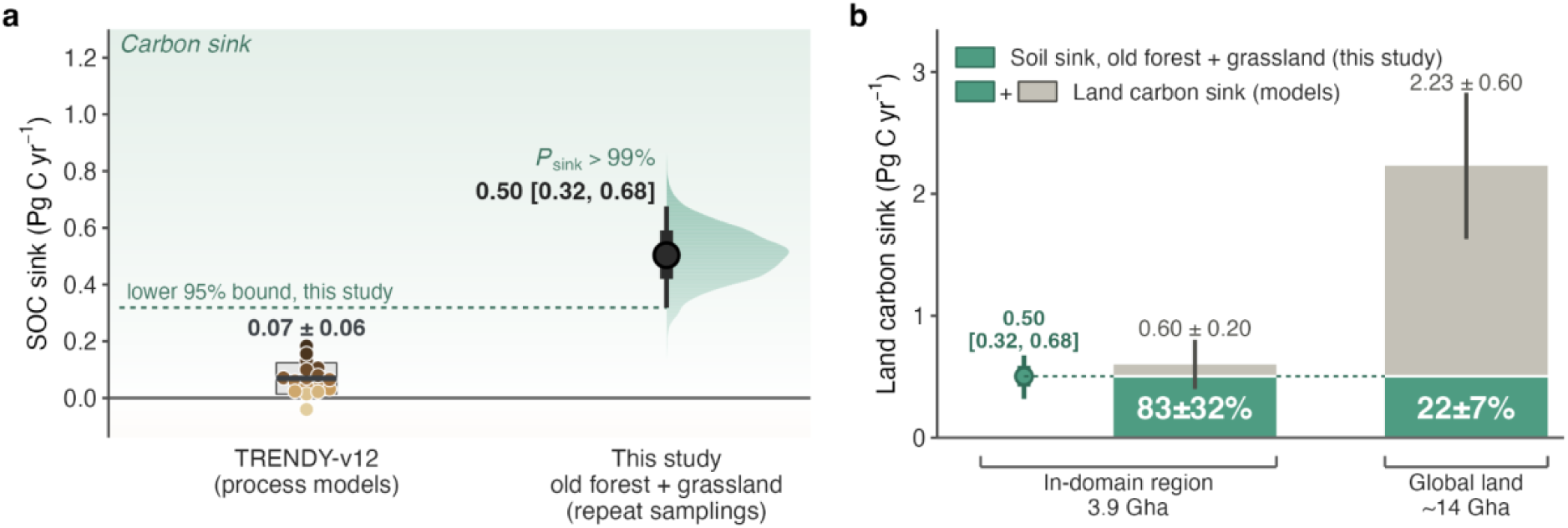
Comparing SOC sink with modeled results. (**a**) Comparison between the observational-based natural SOC sink and TRENDY-v12 (*26*) model estimates over the same spatial and temporal domain (in-domain old forests and grasslands, 1992–2020). On the left, the black horizontal line and gray rectangle represent the TRENDY-v12 multi-model mean and standard deviation (values annotated above). Each circle represents one model estimate, with values reported in Supplementary Table S2. On the right, the half violin indicates the posterior estimate of in-domain SOC sink in old forests and grasslands during 1992–2020 from this study. The black point indicates posterior mean, and the thick and thin bars indicate the 66% and 95% credible intervals. The annotated sink value is the mean and [2.5%, 97.5%] percentile of the posterior estimate. *P*_sink_ indicates the posterior probability that the SOC change is positive. (**b**) Contribution of the in-domain SOC sink to Global Carbon Budget’s (GCB) modeled land sink over the last three decades (*9*). On the left, the green point indicates the posterior mean of the in-domain SOC sink in old forests and grasslands during 1992–2020. The thick and thin bars indicate the 66% and 95% credible intervals. In the middle, the rectangle represents the in-domain modeled land sink during 1992–2020. Rectangle height and error bar (annotated) indicate the multi-model mean and standard deviation among 7 GCB participant models (*29*). On the right, the tall rectangle represents the global modeled land sink. Rectangle height and error bar (annotated) are derived from the mean estimates and uncertainties of GCB global land sinks averaged over the decades of 1990s, 2000s, and 2010s.

To evaluate soils’ role in the land sink, we compared the observation-based SOC sink with the modeled land sink (net carbon gain in vegetation, litter and soil) from the GCB. We restricted the gridded land sink estimates of seven GCB models (29) to the analysis domain, and derived the in-domain land sink as the multi-model mean and standard deviation. Our observational SOC sink accounts for 83±32% of the modeled land sink within this domain (Fig. 4b). This in-domain SOC sink, quantified on 27% of applicable old forest and grassland area, still represents 22±7% of the global modeled land sink (Fig. 4b). This result indicates that soils account for most of the modeled land sink in the area represented by repeated soil measurements. It provides direct observational support for the hypothesis that non-living C pools stored a large portion of the recent land C uptake, which was previously inferred as a residual (*5*).

## Discussion

Soils gained 0.75 [0.50, 1.03] Pg C yr^-1^ during 1992–2020 in the area represented by repeated soil measurements. In old forests and grasslands, this sink is about seven times the sink simulated by process models and equivalent to 83±32% of the modeled land sink. The models therefore assign too little of the land sink to soils in this area, or underestimate the land sink, or both. The partitioning affects model projections of the land sink, because carbon generally resides longer in soils than in vegetation (*14*). However, the change of boreal old forests from a sink to a source in the 2010s indicates that SOC gains can also reverse within a decade.

### I. Potential mechanisms for SOC change

Here we discuss the candidate mechanisms consistent with the patterns of recent SOC change (Fig. 3, Extended Data Tables 2, 3). The SOC accumulation in young forests is consistent with the increase in productivity and in C inputs to soils as trees grow (*6*, *30*, *31*). Temperate old forests lost SOC in the 1990s and gained SOC in the 2000s and 2010s (Fig. 3). The loss in the 1990s is consistent with the SOC loss measured in forests of the German Alps, which was attributed to faster decomposition of particulate organic matter (POM) under warming (*32*, *33*). Boreal old forests gained SOC in the 1990s and 2000s (*34*, *35*). Their sink weakened in each decade, and became a source in the 2010s (Fig. 3, Extended Data Table 2). The SOC gain in boreal old forests before the 2010s is consistent with the increase in plant productivity at high latitudes. Rising CO_2_ and warming have increased growth and lengthened the growing season in boreal regions (*36*, *37*). Warming also increases N mineralization, which increases the N available for plant growth (*38*, *39*). Because low temperatures slow decomposition, the additional inputs may have accumulated mainly as POM (*40*, *41*). The weakening of SOC sink in Russian and Canadian forests was also simulated by models calibrated with local data (*2*). This weakening may be explained by two mutually non-exclusive mechanisms. First, continued warming could enhance heterotrophic respiration (*20*) and POM decomposition (*42*), which might have surpassed C inputs to soils by the 2010s. Second, disturbances have increased in boreal forests, including insect outbreaks, wildfire, and drought (*2*, *43*). Fire removes soil organic matter by combustion, and drought reduces productivity and C inputs to soils. Tree mortality adds dead roots and litter in the short term, but reduces the inputs from living roots until the stand recovers. This shift from sink to source suggests that the response of boreal SOC to warming may have changed sign. Through the 1990s and the 2000s, warming and rising CO_2_ may have increased productivity and SOC. In the 2010s, higher decomposition, disturbance, or both, may have exceeded this gain. Despite this shift, atmospheric inversions report that the land sink in the Northern Hemisphere has been relatively stable during the past three decades (*9*). This contrast suggests that Northern vegetation sink, or SOC sink outside of the analysis domain, increased to compensate for the weakened in-domain SOC sink.

The SOC accumulation in temperate grasslands (Fig. 3, Extended Data Table 2) is consistent with SOC gains under elevated CO_2_ observed in grassland CO_2_ experiments (*19*). Elevated CO_2_ can increase C inputs to grassland soils through roots, root exudates and mycorrhizal fungi, which are efficient sources of SOC (*33*, *44*, *45*). Longer growing seasons under warmer and wetter climates may have increased these inputs further (*46*). Previous regional studies found that grasslands in Europe accumulated SOC (*47*), but grasslands in Northern China experienced SOC losses that cannot be attributed to grazing (*48*). Qualitatively consistent with these patterns, we found grassland SOC accumulation rates (Mg C ha^-1^ yr^-1^) to be higher in Europe than in Asia (Extended Data Table 3). This difference may reflect the higher N deposition in Europe, which increases grassland productivity (*49*) and therefore C inputs to soils.

### II. Limitations and future research priorities

This analysis is subject to several limitations inherent to soil data syntheses. First, we evaluated SOC changes only at the top 30 cm of mineral soil due to limited data on deeper soils. The role of deeper soils and the litter layer in land C uptake requires further investigation (*50*). Second, although this analysis focused on natural SOC changes by excluding sites with active land-use change, there are not sufficient data to explicitly partition SOC changes caused by prior land use history, concurrent management practices (e.g., selective harvesting, grazing), and disturbance history (e.g., fire, windthrow, insect outbreaks). Detailed site history records on every repeatedly sampled site would improve future SOC sink assessments and partition the cause of SOC change among background climate change, disturbance, and management. Third, despite their unique biogeochemical processes (*51*), wetland and permafrost soils were not treated as explicit categories because the database is limited in sites that are known to be waterlogged. Only around half of the sites report information on waterlogging, within which only 32 sites are known to be waterlogged. These soil systems should be further studied with specialized data and tools.

Fourth, SOC varies at fine scales (<1 m^2^), which makes SOC change difficult to detect at a single site (*52*). We accounted for this heterogeneity as statistical variance through random effects and large-sample aggregation. Our estimates therefore apply to broader spatial scales rather than individual sites.

This synthesis reveals that although the research community has accumulated a rich set of repeated soil samplings in North America and Europe over the past five decades, accessible repeated samplings remain scarce beyond these regions. Particularly, the boreal region in Northern Russia and global tropical ecosystems have almost no repeated measurements. With the existing data, we can estimate the SOC sink for 30% of the area of forests and grasslands without land-use change. Three actions would extend this analysis domain: resampling sites that were sampled once in data-scarce regions, making more existing repeated measurements publicly accessible (*53*, *54*), and standardizing sampling protocols and data formats (*55*).

## Conclusion

We provide the first observation-based estimate of the natural SOC sink, from repeated soil sampling at 4,883 forest and grassland sites. In the area represented by these sites, 30% of the global area of forests and grasslands without land-use change, soils gained 0.75 [0.50, 1.03] Pg C yr^-1^ from 1992 to 2020. In old forests and grasslands, this sink is about seven times the sink simulated by process models. The sink weakened in the 2010s, when soils in boreal old forests changed from a sink to a source, so its persistence is uncertain. These estimates can be used to evaluate SOC change in process models and in the Global Carbon Budget.

## Methods

### Data compilation

We compiled repeated soil samplings in forests and grasslands from databases (*56–65*) (Supplementary Table S3), published literature (*32*, *48*, *66–82*), and unpublished data contributed by the authors. To synthesize data from the literature, we conducted a Web of Science search with the string “TOPIC: (change* NEAR “soil carbon”) OR (change* NEAR “soil organic” matter) OR (soil carbon organic matter change gain loss) OR (soil carbon organic matter long-term) sampl*” to include repeated soil samplings and “NOT TOPIC: (agricultur* OR crop* OR fallow* OR till* OR cultivat* OR farm* OR graz* OR rotation OR paddy OR yield OR conversion OR plantation OR infrared OR infra-red OR spectroscopy OR dissolv* OR chronosequence OR incubat*)” to rule out cultivation, indirect measurements, and incubation experiments. The search captured very few relevant studies, likely because repeated measurements from control plots are often embedded in studies focused on experimental manipulations or treatments and are not explicitly labeled as repeated soil samplings. The criteria for data collection were the following. (1) Coordinates must be reported or can be found by site name. Land cover of the site must be reported by the data source. (2) Field measurements of soil organic carbon (SOC), soil organic matter (SOM), or total carbon (TC) in concentration or stock must be reported at least twice in different years on the same spatial unit defined by the source.

All observations in each soil carbon time series (repeated measurements) must report the same variable (all SOC concentration or all SOM stock, etc.) with identical sampling and analysis protocols. Upper and lower depths must be reported for all soil layers. (3) Sites included must have no known record of cultivation or experimental manipulations (e.g., fertilization, fencing) during the sampling period. Consistent with the study goal of evaluating SOC changes in forests and grasslands without land-use and land-cover change, we excluded sites where land cover type was changed during the sampling period. We included sites with other management and disturbances, such as domestic animal grazing, prescribed fires imitating natural frequency in grasslands, pest infestations including beetles, and single-selection harvesting in forests. (4) Data must come from sources with documented sampling and analysis protocols: inventories and observation networks, published studies, or datasets contributed by authors. In the end, data from 6,739 plots in 6,074 sites were collected. Each plot had been observed for at least two separate years, forming a time series. Some sites had multiple plot-level time series if more than one replicate plot was sampled repeatedly.

### Processing soil carbon data

Before statistical analysis, we processed the raw data of SOC measurements into the top 30 cm SOC stocks. We collected values of soil bulk density (BD) and soil coarse fragments from data sources when available and gap-filled the rest using the fine earth BD and volumetric coarse fragment fraction estimates from SoilGrids 2.0 (*83*). Soil layers that reported SOC concentrations were first converted to SOC stocks using the following formulas:

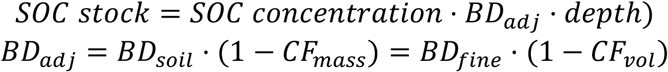

where *BD_adj_* is the mass of fine earth (all soil particles < 2 mm) per unit volume of bulk soil, which was calculated from either *BD_soil_* or *BD_fine_* based on the definition of BD reported from data source. *BD_soil_* is the density of bulk soil, including the mass of rocks and coarse fragments. *BD_fine_* is the mass of fine earth per volume of fine earth, excluding coarse fragments. *CF_mass_* is the coarse fragment fraction by mass, and *CF_vol_* is the volumetric fraction of coarse

fragments. *BD_fine_* and *CF_vol_* data were gap-filled by SoilGrids 2.0 (*83*). The threshold between fine earth and coarse fragments was defined as 2mm in both SoilGrids 2.0 (*83*) and data sources. When needed, *CF_mass_* and *CF_vol_* were converted between each other assuming a coarse fragment density of 2.6 g/cm^3^ following the equations in (*84*). Within a SOC time series, if BD was reported for one visit but not another, the reported BD was used to calculate soil C stock for other visits of the plot for consistency, to avoid spurious SOC change driven by the difference between gap-filled and field-based BD of the same plot. Sixty-two plots within 43 sites reported soil organic matter (SOM) concentration, where we used the conversion factor *SOC* = 0.5 ⋅ *SOM* (*85*). Seventeen plots within 6 sites reported TC stock or concentration. We took these values as a proxy of SOC stock or concentration in data processing and analysis.

After converting all layered measurements into SOC stocks, we adjusted layered SOC stocks within each soil profile into cumulative SOC stock at the top 30 cm following the methods in (*86*). For each soil profile with at least two layers reported, we fitted a log-log curve of cumulative SOC stock as a function of lower depths, and then predicted the top 30 cm SOC stock from the fitted curve. We applied a threshold of R^2^ ≥ 0.8 to ensure reliability of the curves and excluded plots where soil profiles had poor fits. For single-layered soil profiles, we estimated the top 30 cm SOC stocks using biome-specific slope coefficients from empirical estimates in (*86*).

To clean the data before analysis, we removed observations that were outliers in terms of their spatiotemporal representation in the dataset or showed extreme values that could represent errors. One observation before 1900 was excluded, making all of the time series span 1957– 2022. To exclude extreme values in SOC changes and potential data entry errors, we calculated the ratios between the first and repeated measurements at each plot. The resulting ratios have a right-skewed distribution. We took the natural logarithm of this ratio and excluded time series whose logged ratios were beyond two standard deviations from the mean. Finally, 5,364 repeatedly sampled plots in 4,883 forest and grassland sites were included in the analysis (Fig. 1). Around 90% of the time series span five years or longer. Plots were revisited a median of 6 years apart over a median span of 9 years, and a quarter of all plots (1,339 plots in 1277 sites) were sampled three or more times, up to 19 years.

Out of the 5,364 plots (from 4,883 sites), 2,711 plots (2,291 sites) are not waterlogged, 42 plots (32 sites) are waterlogged, and 2,611 plots (2564 sites) have no records of waterlogging. Disturbances were recorded in 364 plots (226 sites). Another 386 plots (252 sites) were monitored for disturbance but reported no events. The rest of 4,614 plots (4,408 sites) were not monitored for disturbance. Out of all grassland sites (2,590 plots from 2,510 sites), 2,362 plots (2,346 sites) were grazed, 9 plots (7 sites) were managed in other forms, 122 plots (114 sites) were unmanaged, 97 plots (45 sites) do not have management information. Out of all forest sites (2,774 plots from 2,376 sites), 642 plots (631 sites) were managed, 400 plots (314 sites) were unmanaged, 1,732 plots (1,432 sites) do not have management information. Some sites contain replicate plots with different land cover, waterlogging, management, and disturbance records, so the site counts can sum to more than 4,883.

### Dividing young and old forests

We collected forest stand age and successional stage from data sources and gap-filled the missing values with the World Forest Age Map (*87*). A forest plot was grouped as young forest if the source reported an earlier successional stage than a mature or old-growth forest (usually reported as early, middle, or late successional stage), or if the stand age was less than 60 years in the year of first SOC measurement. A forest plot was grouped as old forest if the source reports the plot as a mature or old-growth forest, or if the stand age was greater than or equal to 60 years in the year of first SOC measurement. In the database, the SOC rates of change in young forests are significantly higher than old forests, supported by data with (p=0.002) and without (p=0.03) forest plots whose stand ages were gap-filled (Wilcoxon signed-rank test, H_1_: young forest > old forest). This result is qualitatively consistent with the observed net primary productivity (NPP) plateau and decline as forest stands grow after 40–60 years (*88*), potentially causing SOC accumulation rates to decrease shortly after the productivity plateau (*89*).

### Statistical analyses

We fitted a Bayesian generalized additive mixed model (GAMM) to estimate SOC stocks in different years, land cover types, climate conditions, and soil properties using the R package *brms* (*24*). The dependent variable was log-transformed top 30 cm SOC stock in a given year.

The independent variables included land cover type (young forest, old forest, grassland), mean annual temperature (MAT) and mean annual precipitation (MAP) in the year of soil sampling, soil N content, soil clay content, and soil pH. Numeric variables were all standardized before fitting the model. MAT and MAP were extracted from the Climatic Research Unit gridded Time Series (CRUTS v4.07) (*90*). We chose the mean temperature and precipitation of the sampling year as a parsimonious approximation of the prevailing climate state relevant for SOC dynamics at each site. This choice avoids introducing an arbitrary, globally uniform memory window while still capturing broad spatial gradients and long-term differences in climate that drive SOC inputs and losses. Soil N content, clay content, and pH were collected from source data where available or gapfilled from SoilGrids 2.0 (*83*). MAT and MAP vary with the sampling year, whereas land cover type and soil properties (N content, clay content and pH) have one value per plot that is constant over time. Soil N content changes over time together with SOC because of the stoichiometry of soil organic matter. We used a time-invariant soil N value so that this predictor represents differences in nutrient status and background SOC storage among sites, and not changes in N availability within a site. Because soil N and SOC are correlated, we tested whether the estimated SOC sink depends on this predictor by refitting the model without soil N. The SOC sink within the analysis domain for 1992–2020 was 0.58 [0.33, 0.85] Pg C yr^-1^ without soil N and 0.75 [0.50, 1.03] Pg C yr^-1^ in the main model, and the 95% credible intervals overlap. The model without soil N had lower predictive accuracy (difference in expected log predictive density, -130.3 ± 20.6; ref. *91*), so we retained soil N in the main model.

We used a nonlinear model structure to capture the potential nonlinear (curvilinear) temporal SOC trends under different conditions. We allowed MAT and MAP to influence SOC temporal trends nonlinearly by fitting two tensor product smooths of year and MAT, year and MAP, respectively (Supplementary Fig. S3, Supplementary Table S2). To capture different SOC responses in different land covers, the coefficients in these smooth terms were fitted independently for each land cover type (young forest, old forest, grassland). We specified a first-order penalty on these smooths for more conservative predictions on SOC changes. We allowed soil N content, clay content, and pH to influence the slopes of SOC temporal trends, assuming their effects were consistent across all land cover types. Land cover type was included as categorical predictors to account for different background levels of SOC storage in grasslands, young forests, and old forests. Random intercepts at the site and plot level were included to account for SOC variations across space. We did not include grazing status to grasslands for two reasons. First, grazing status is not separable from land cover given the database: most grasslands sites were domestically grazed and most forests were not. Second, we could not assign grazing status accurately to the prediction domain, which would be necessary during upscaling if the model includes this term.

We used vague priors *N*(0,5) for the fixed effects and followed the defaults of the *brms* package (*24*) for the priors of random effects and smooth terms. Three Markov chains were run for 26,000 interactions individually, with the first 1,000 iterations as warm up and a thinning interval of 50. The model produced 500 posterior samples from each chain, 1,500 posterior samples in total. All parameters had R-hat values below 1.05. The Markov chain Monte Carlo (MCMC) chains mixed adequately based on visual evaluation of trace plots. No divergent transitions were detected during sampling. Energy Bayesian Fraction of Missing Information (E-BFMI) indicated adequate exploration of the posterior distribution, suggesting that the Markov chains sampled efficiently. Posterior predictive check shows no warning signs. Bulk and tail effective sample sizes were generally above 300 across parameters, indicating adequate sampling efficiency and well-estimated posterior distributions. Based on diagnostics above, we determined that the model is converged and the posteriors are reliable for inference. The conditional Bayesian R-squared (*25*), which includes the variance of random effects, is 0.90 ± 0.0012. The marginal Bayesian R-squared (*25*), which excludes the variance of random effects, is 0.49 ± 0.008. We interpreted the 2.5% and 97.5% percentile of the posterior predictive distributions as 95% credible intervals of the estimates, and calculated standard errors of the estimates as the standard deviation of the posterior predictions.

### Upscaling

We used the fitted model to estimate regional and total natural SOC changes by applying the model predictions on areas of forests and grasslands without land cover change during 1992– 2020, based on the yearly European Space Agency’s (ESA) Climate Change Initiative (CCI) land cover maps (*92*). The data products used in gap-fillings from CRUTS (*90*) and SoilGrids 2.0 (*83*) were used again for prediction (Supplementary Fig. S4). Raster layers of soil N content, soil clay content, and soil pH were resampled to match the coarser 0.5-degree resolution of MAT and MAP layers from CRUTS.

To avoid over-extrapolation, we upscale SOC changes only to the 0.5-degree pixels well-represented by the input data. We consider a pixel to be well-represented by the input data (in-domain) if its predictor values lie within the input data’s predictor value ranges without outliers. The estimated area extent is shown in Supplementary Fig. S1 and Extended Data Table 1.

For each land cover type, we predicted the 0.5-degree pixel-level SOC stocks (Mg C ha^-1^) in the year 1992, 2000, 2010, and 2020 using historical MAT and MAP data of the corresponding years. The estimation of the SOC sink started in 1992 because it was the earliest year with an available land cover map. Then, for each decade of interest, the rate of SOC change per area (Mg C ha^-1^ yr^-1^) at the pixel level for each given land cover type *ΔSOC_p_*_,*lc*_ was calculated by taking the difference between the predicted SOC stock in the initial year (yr0) and the last year (yr1), normalized by the time duration.

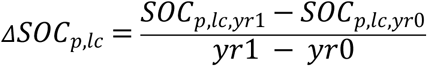

The regional and total SOC stock changes (Tg C yr^-1^ or Pg C yr^-1^) for each land cover were calculated by summing pixel-level SOC changes multiplied by the applicable area in each pixel.

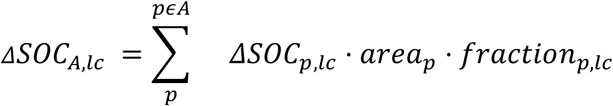

where *ΔSOC_A_*_,*lc*_ is the estimated annual SOC stock change in a given region *A* for a given land cover type *lc* (young forest, old forest, or grassland). *ΔSOC_p_*_,*lc*_ is the pixel-level rate of SOC change per area for a given land cover. *A* is the region of interest. *area_p_* is the cell size of the given 0.5-degree pixel. *fraction_p_*_,*lc*_ is the area fraction of a given land cover without land cover class change during 1992–2020 in a 0.5-degree pixel.

We applied the same land cover fractions (*fraction_p_*_,*lc*_) for different decades, so the differences in decadal SOC changes only stem from the SOC change rate and not from changes in predicted area (Extended Data Table 1). We calculated the area fractions using the following method. First, we downloaded the yearly ESA CCI land cover maps (*92*) for years 1992, 2000, 2010, and 2020 at the resolution of 300 meters. For each year, we classified the 300-m pixels as forest if the pixels had the land cover class of “Tree cover” or “Mosaic Tree and shrub (>50%)” (land cover classes 50-100), and as grassland if they had the land cover class of “Mosaic herbaceous cover (>50%)” and “Grassland” (land cover classes 110 and 130). To filter for areas without land cover change, we filtered pixels by their values in “change count”, which records the number of land cover changes of a given pixel from 1992 to the year of the map. To calculate the grassland fraction for each 0.5-degree pixel without land cover change during 1992-2020, we filter the pixels that are assigned as grassland in the 2020 land cover map to have a “change count” class equal to 0, meaning that the pixels’ land cover class has not changed any year from 1992 to 2020. The grassland fraction for each 0.5-degree pixel, *fraction_p_*_,*lc*=*grassland*_, was calculated with zonal statistics as the areal fraction of all 300-m grassland pixels without land cover change in the 0.5-degree pixel *p*.

To assign the fractions of young and old forests during 1992–2020, we downloaded the World Forest Age Map (*87*) and resampled the 1-km-resolution map to match the 300-m resolution of the CCI land cover maps. As forest stands age during the past three decades, young forests would transition into old forests in a pixel, resulting in old forest fractions accumulating and young forest fractions decreasing over time. To account for forest aging, we calculate the fractions of young and old forests during 1992 and 2020 by averaging the young and old forest fractions calculated from three separate decades. We first find forest pixels in the CCI map without land cover change during each decade (1992–2000, 2000–2010, 2010–2020). For example, for decade 2000-2010, we filtered the forest pixels in the 2000 map that have the same “count change” value in the 2000 map and the 2010 map, meaning there is no additional land cover change on that pixel during 2000 and 2010. Following this method, we produced one 300-m resolution raster layer representing forests without land-cover change for each decade of 1992–2000, 2000–2010, 2010–2020. For each decadal layer, we assigned the 300-m forest pixels into young and old forest using the resampled World Forest Age Map, which estimates forest age at the year 2010. First, we assigned forest age to each pixel at the beginning of each decade. For example, for decade 2000-2010, if a pixel is estimated to have age 55 in 2010 by the World Forest Age Map, we assume that the pixel would have forest age of 45 in the year 2000. Then, we classified the pixels in each decadal layer into young forest (<60 years) or old forest (>=60 years) based on their forest ages at the beginning of the decade. After acquiring the decadal layers containing young and old forest pixels of 300m resolution, we calculated fractions of young and old forests by running zonal statistics of all young and old forest 300-m pixels in each 0.5-degree pixel. This calculation gives us estimates of young and old forest fractions for each decade (E.g., *fraction_p_*_,*young forest*,1992−2000_). Finally, we calculate young and old forest fractions during 1992-2020 by averaging their fractions across all three decades.

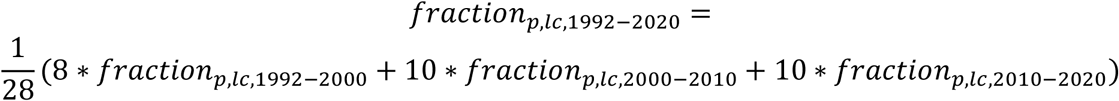

The SOC change at pixel levels was predicted for all 1,500 sets of posterior samples and summed into regional and global extents, yielding posterior predictive distributions of pixel-level SOC change, regional SOC sink, and total SOC sink. The standard errors presented in pixel-level SOC change (Extended Data Figs. 1-5), regional and total SOC sink (Extended Data Table 2), and region-average SOC change rate (Extended Data Table 3) were calculated as the standard deviation of their corresponding posterior predictive distributions. The 66% and 95% credible intervals of regional and total SOC sink estimates were drawn from their posterior predictive distributions as the 17%–83% percentile and 2.5%–97.5% percentile range, respectively.

Strengthening and weakening of the SOC sink between two decades were calculated as the SOC sink (source) at the later decade minus the SOC sink (source) at the earlier decade. This calculation was also made for all 1,500 posterior samples to generate posterior predictive distributions. Extended Data Figs. 6 and 7 show the mean magnitudes of pixel-level SOC sink change and posterior probability of the changes. In Fig. 2 and Extended Data Table 2, if more than 95% of the posterior estimates show the same trend in SOC sink change between two decades, we consider the SOC sink to have strengthened or weakened detectably.

### Comparison with TRENDY models

We compared the observation-based estimate of the SOC sink in old forests and grasslands to models from the TRENDY-v12 model ensemble (*26*). To exclude the impact of land-use and land-use change, we rely on the S2 simulation, which includes transient atmospheric CO_2_, transient climate, and fixed pre-industrial land-use (*26*). We excluded young forest SOC sink from the comparisons because the TRENDY-v12 S2 simulations are not designed to represent young forest recovery. We calculated the modeled SOC sink as changes in the cSoil variable, excluding the litter (cLitter) and coarse woody debris (cCwd) variables from each model. CLM5.0 is excluded from the comparison because its cSoil inherently includes cLitter. Similarly, CABLE-POP was excluded because its cSoil includes cCwd.

First, we calculated the models’ total SOC stocks in 1992 and 2020 for in-domain old forests and grasslands (Supplementary Fig. S5), keeping the spatial and temporal span consistent with this analysis. Because the TRENDY S2 simulation used fixed pre-industrial land cover, we multiplied the pixel areas with the land cover fractions used in this study. Then, we calculated the SOC sink for each model as the average annual change in total SOC stock between 1992 and 2020 (Supplementary Table S2). Finally, we presented the ensemble mean and standard deviation of the modeled in-domain SOC sinks (Fig. 4a). One distinction between this comparison is that the TRENDY models include C changes from soils deeper than 30 cm, whereas the observation-based analysis focuses only on the top 30 cm because of data limitations.

### Comparison with the modeled land sink

In the Global Carbon Budget (GCB) (*9*), the land sink (S_LAND_) is defined as the gross increase in terrestrial C stocks driven by global change factors without the direct influence of land use and land-use changes. Therefore, we used the SOC sink estimates of old forests and grasslands to compare with the GCB modeled land sink estimates. The young forest was excluded because the C gain from recovering young forests is classified as land use change emissions rather than part of the land sink.

We masked the gridded S_LAND_ fluxes from O’Sullivan and authors (*29*) to derive S_LAND_ in the spatial domain of the SOC sink estimates (in-domain S_LAND_) (Fig. 4b). This study reported gridded land fluxes for seven TRENDY models corrected for the Replaced Sinks and Sources (RSS) bias (*29*). The same corrected models were used in the GCB report for regional and spatial S_LAND_ estimates (*9*). Because the gridded S_LAND_ fluxes with RSS correction already pre-adjusted pixel-level fluxes to account for realistic land cover fractions, we do not further apply the land cover fractions from this study. After calculating the 1992–2020 mean of gridded fluxes for each model, we masked the gridded layers to include only the 0.5-degree pixels we consider in-domain for old forests and grasslands. We derived in-domain S_LAND_ by first calculating the total C fluxes for each pixel (pixel-level fluxes in kg C m^-2^ yr^-1^ times corresponding pixel area), and then summing the total C fluxes for all in-domain pixels.

For the global S_LAND_ in Fig. 4b, we averaged the GCB-reported S_LAND_ values over the 1990s, 2000s, and 2010s. The GCB modeled S_LAND_ was derived from the same 7 models with RSS correction from O’Sullivan and authors (*29*), and the remaining 15 TRENDY models adjusted with empirical reductions as RSS correction, which was estimated from SLAND∼RSS relationships in the 7 models.

The percentages of the SOC sink in in-domain S_LAND_ (83±32%) and global S_LAND_ (22±7%) are calculated as the SOC sink estimate divided by the land sink (Fig. 4b). Uncertainty is propagated assuming the estimated SOC sink and land sink are independent random variables, following:

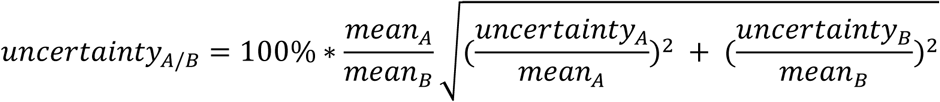

where A (numerator) is the in-domain SOC sink estimate in old forests and grasslands (0.5±0.07 Pg C yr^-1^), with uncertainty defined as the standard deviation of its posterior distribution. B (denominator) is either the in-domain land sink (0.60±0.20 Pg C yr^-1^) or global land sink (2.23±0.60 Pg C yr^-1^). The uncertainty of the in-domain land sink is the standard deviation of estimates from the 7 models in (*29*). The uncertainty of the global land sink is the averaged uncertainty values for the GCB global land sink over the 1990s, 2000s, and 2010s (*9*).

We acknowledge a few technical distinctions between the in-domain SOC sink in old forests and grasslands and the definition of S_LAND_ in the GCB. First, the GCB defines S_LAND_ as the C accumulation of terrestrial ecosystems that is driven by global change factors including CO_2_, N deposition, and climate change, while land use change emission (ELUC) is defined separately as the C changes in terrestrial system caused by land-use, land-use change, and forestry (*9*). Together, the C accumulations in S_LAND_ and C loss in ELUC add up to the net land flux (NetLAND). Excluding young forests, which undergo recovery and whose C fluxes belong to ELUC, brings the SOC sink estimate much closer to the GCB definition of S_LAND_. Applying land cover fractions without land-cover changes further excluded C changes defined as ELUC from the SOC sink estimate. However, such land cover fractions also excluded areas that have experienced land-cover class changes because of CO_2_, N deposition, and climate (greening, woody encroachment, natural disturbances), and SOC change in such areas would be included in S_LAND_ per GCB definitions. Nevertheless, this distinction would not lead to an overestimated observation-based SOC sink over the analysis domain, because it makes the upscaled area more conservative than the GCB definition. Second, limited by data, this study does not explicitly partition climate-induced SOC changes from SOC changes caused by management or human activities, as they are separately defined in GCB. Future observation-based studies should further partition the causes of SOC changes in natural systems to better benchmark land C fluxes in the GCB.

### Use of Large Language Models

Generative AI (Claude, Anthropic) was used to help write the code that plots the figures. These codes were then manually verified and modified by the authors.

## Data availability

All data will be posted to an online repository upon acceptance. Please contact Ruofei Jia for additional material requests.

## Code availability

All code is available at https://github.com/RuofeiJia/SOCTimeSeries and https://github.com/RuofeiJia/SOC_change_TRENDYv12.

## Acknowledgments

We thank H. Vallicrosa, L. Mirzagholi, S. Bell, Y. Feng, N. Chen, S. Ren, J. Yu, S. Cerasoli, J. Liu, S. Chen for the comments and support. We thank H. Zhu for her contribution in early-stage data exploration. We thank the authors whose published data was included in this analysis. We thank the databases for coordinating and sharing data included in this analysis. We thank the TRENDY team for the provision of the model simulations. This material is based upon work supported by the National Science Foundation under Grant No. DEB-2339051. This research was supported by a seed award from the MIT Climate and Sustainability Consortium. This is a contribution of the MIT Terrer Lab. This work is supported by National Science Foundation under Grant No. DEB-233905. RJ was supported by the Jack Tang Fellowship and Martin fellowship at MIT. AM was supported by a Laboratory Directed Research and Development Program at PNNL. This work was generated using data from the Nutrient Network (http://www.nutnet.org) experiment, funded at the site-scale by individual researchers.

Coordination and data management have been supported by funding to ETB and EWS from the National Science Foundation Research Coordination Network (NSF-DEB-1042132) and Long Term Ecological Research (NSF-DEB-1831944 and NSF-DEB-2425352) programs, and the Institute on the Environment (DG-0001-13). We also thank the Minnesota Supercomputer Institute for hosting project data. Soil analyses were supported, in part, by USDA-ARS grant 58-3098-7-007 to ETB. The evaluation was based on data that was collected by partners of the official UNECE ICP Forests Network (http://icp-forests.net/contributors). Part of the data was co-financed by the European Commission (Data achieved at 10/12/2023).

## Author contributions

R.J. and C.T. designed the study. R.J., C.T., E.F., and J.D. developed the methods in statistical analysis. R.J. and A.M. developed the database format. R.J. and M.S. compiled the database with contributions from G.P., B.B., R.A., N.F., M.B., Y.Z., T.W.B., J.A.D., S.K., K.P., N.E., P.L.P., A.S.M., S.A.P., S.C.R., P.M., S.H., E.T.B., E.W.S., H.B., and A.E. E.T.B. and E.W.S. curated and managed the Nutrient Network data, which were compiled into the database by R.J. Y.M.B. and R.J. conducted the model comparisons. S.S., M.O., P.F. provided data for the TRENDY models and improved the model comparison section. R.J. conducted the analysis and wrote the original draft. C.T., E.F., A.M., Y.M.B., B.B.L., G.P., R.B.J., B.A.H., S.S., M.O., P.F. provided comments that improved the methodology. R.J., C.T., and E.F. revised the draft. All authors contributed to multiple rounds of revisions and the final version.

---

**Extended Data Fig. 1.**
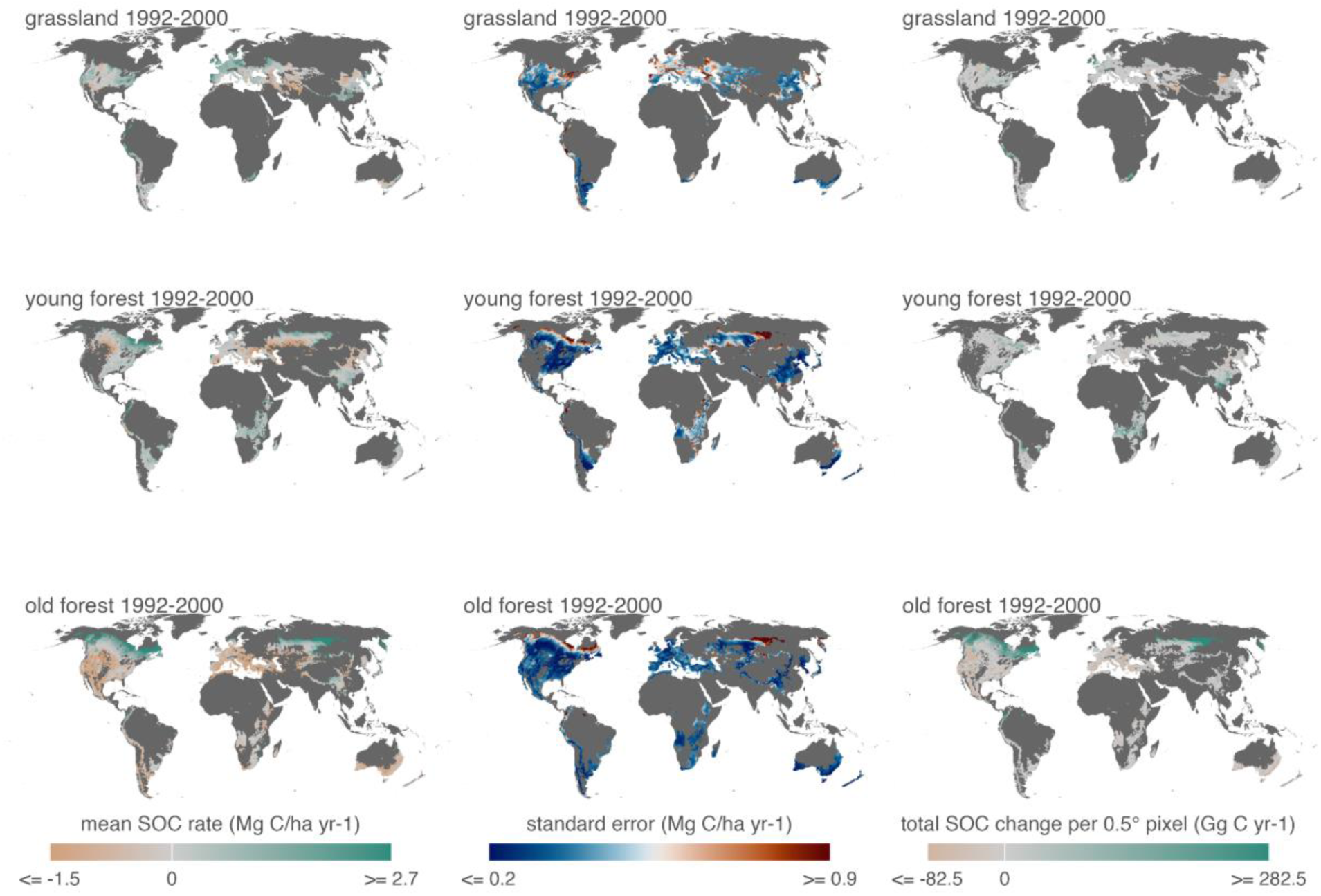
Maps of predicted pixel-level SOC rates during 1992–2000 (column 1), associated standard errors (column 2), and total SOC changes per pixel (column 3). Pixel-level SOC rates were predicted as the posterior means. Standard errors of the SOC rates were calculated as the standard deviations of the posterior predictive distributions. Total SOC changes per pixel were calculated by multiplying the pixel-level SOC rates with each pixel’s area and land cover fraction (Supplementary Fig. S5).

**Extended Data Fig. 2.**
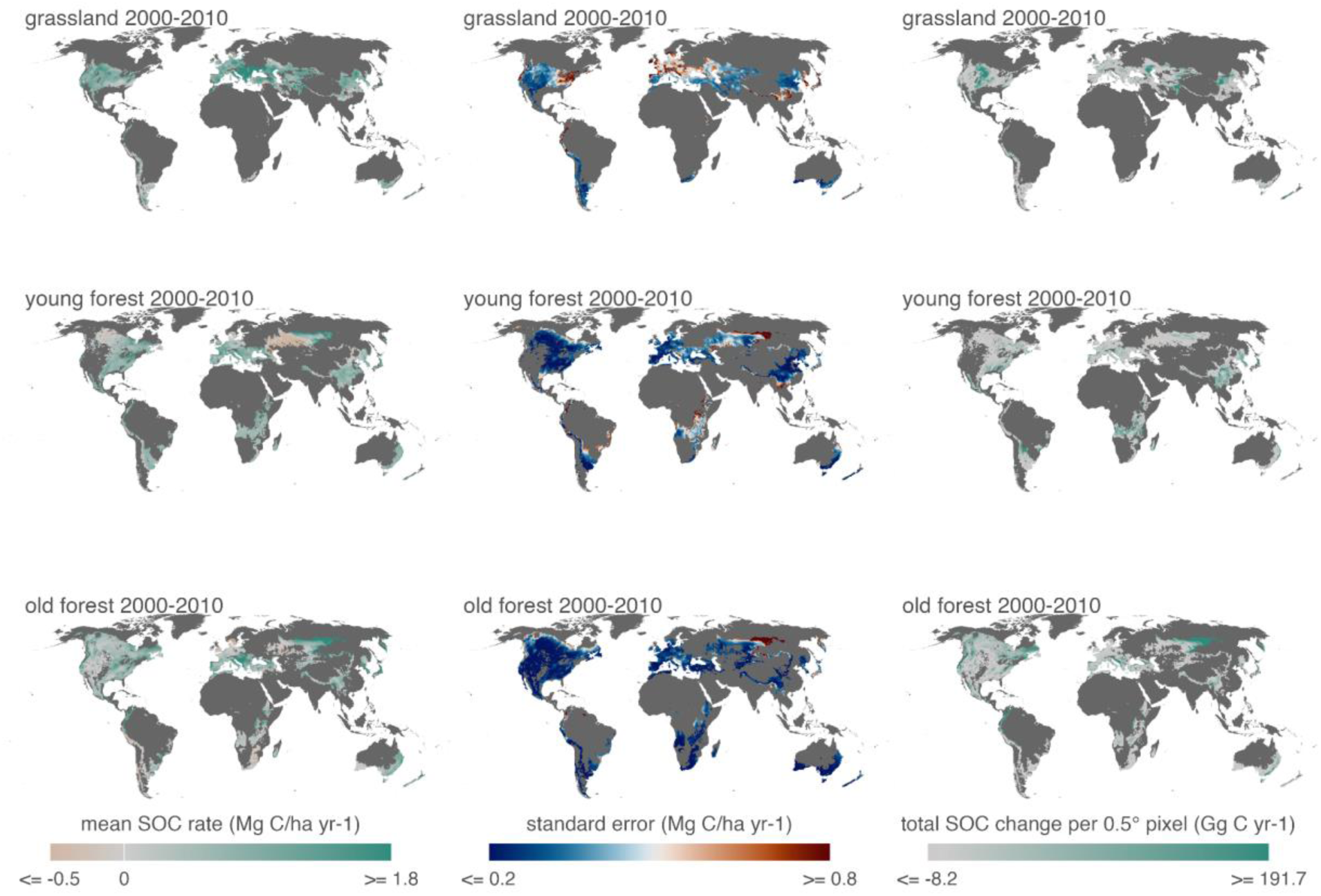
Maps of predicted pixel-level SOC rates during 2000–2010 (column 1), associated standard errors (column 2), and total SOC changes per pixel (column 3). Pixel-level SOC rates were predicted as the posterior means. Standard errors of the SOC rates were calculated as the standard deviations of the posterior predictive distributions. Total SOC changes per pixel were calculated by multiplying the pixel-level SOC rates with each pixel’s area and land cover fraction (Supplementary Fig. S5).

**Extended Data Fig. 3.**
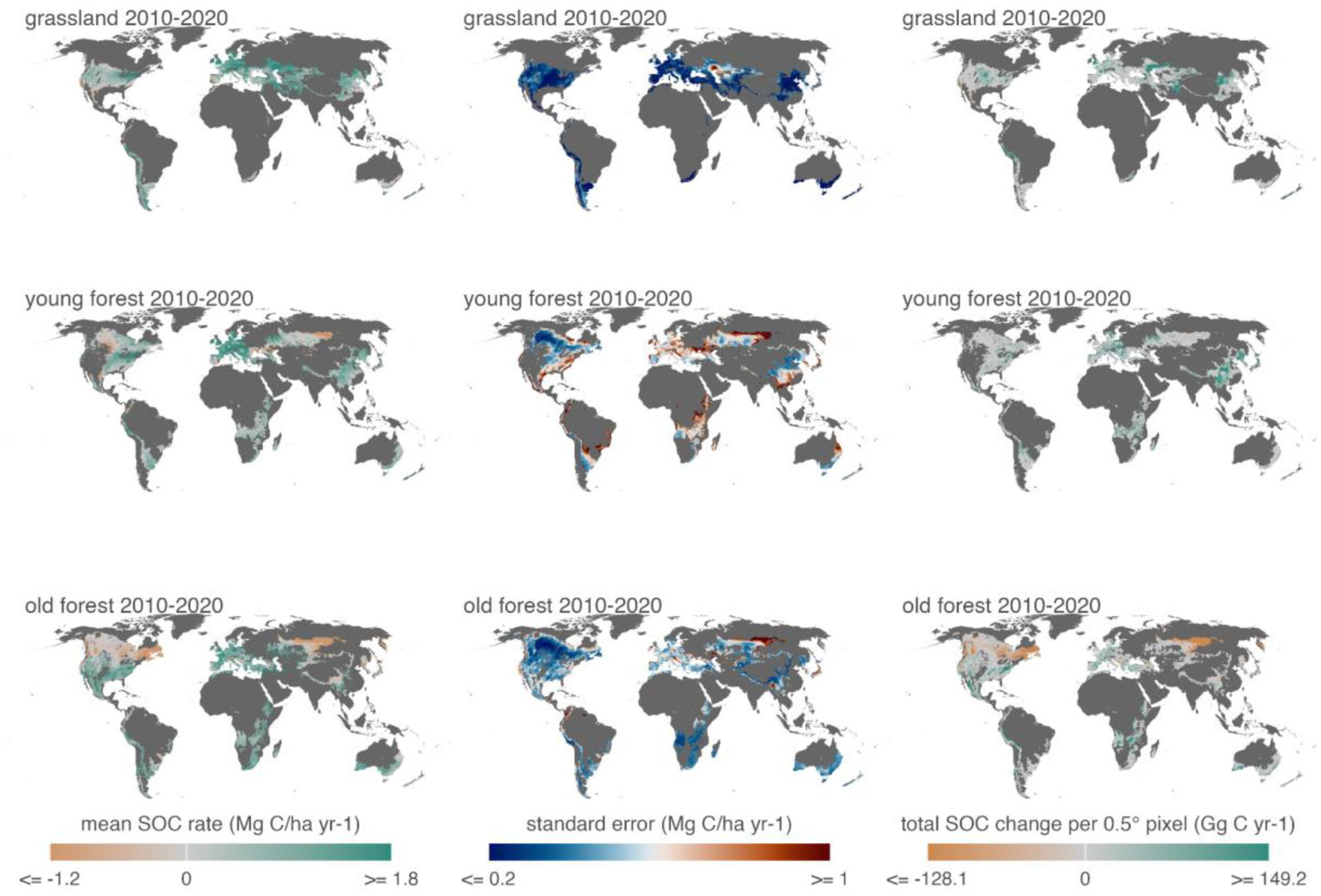
Maps of predicted pixel-level SOC rates during 2010–2020 (column 1), associated standard errors (column 2), and total SOC changes per pixel (column 3). Pixel-level SOC rates were predicted as the posterior means. Standard errors of the SOC rates were calculated as the standard deviations of the posterior predictive distributions. Total SOC changes per pixel were calculated by multiplying the pixel-level SOC rates with each pixel’s area and land cover fraction (Supplementary Fig. S5).

**Extended Data Fig. 4.**
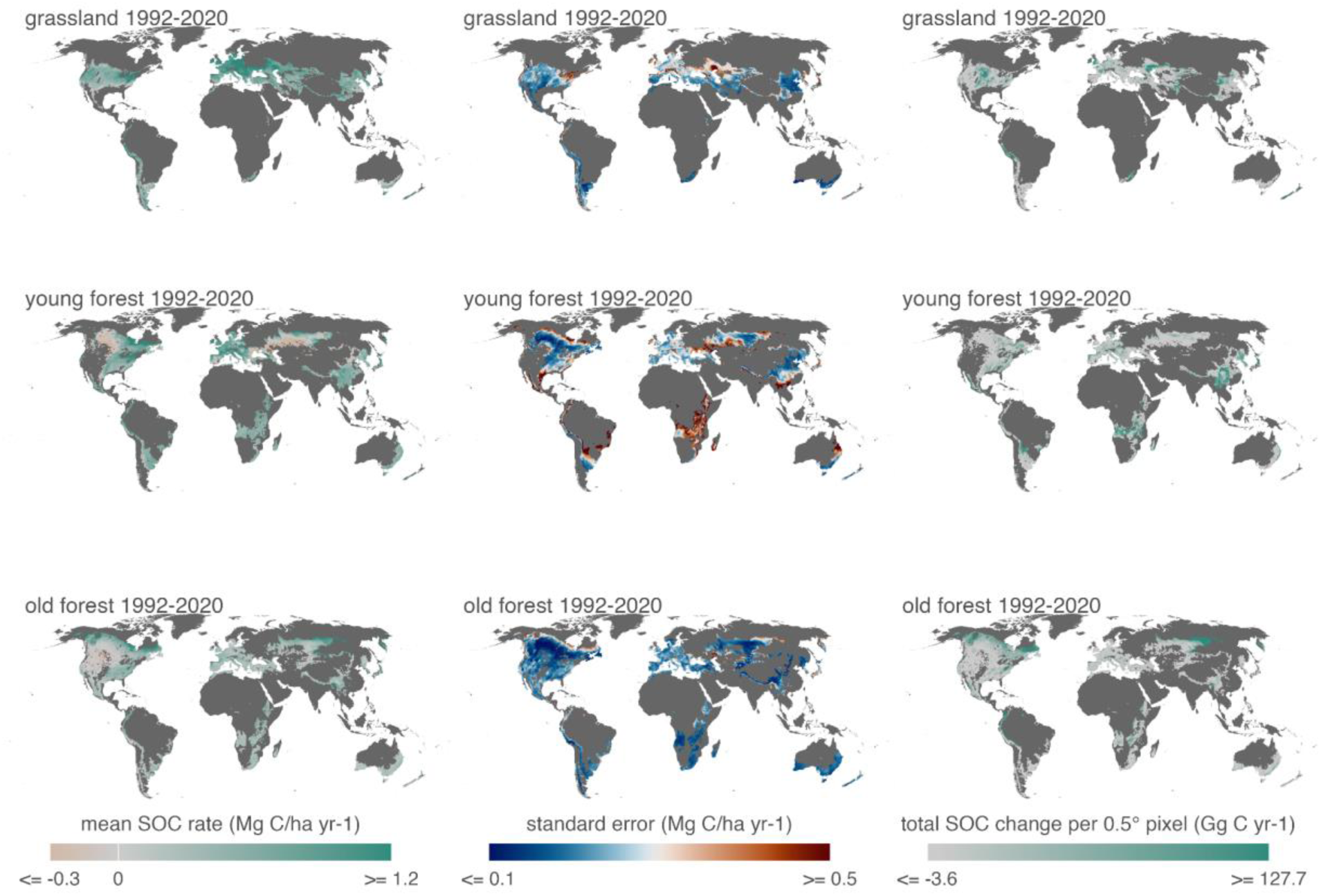
Maps of predicted pixel-level SOC rates during all three decades from 1992 to 2020 (column 1), associated standard errors (column 2), and total SOC changes per pixel (column 3). Pixel-level SOC rates were predicted as the posterior means. Standard errors of the SOC rates were calculated as the standard deviations of the posterior predictive distributions. Total SOC changes per pixel were calculated by multiplying the pixel-level SOC rates with each pixel’s area and land cover fraction (Supplementary Fig. S5).

**Extended Data Fig. 5.**
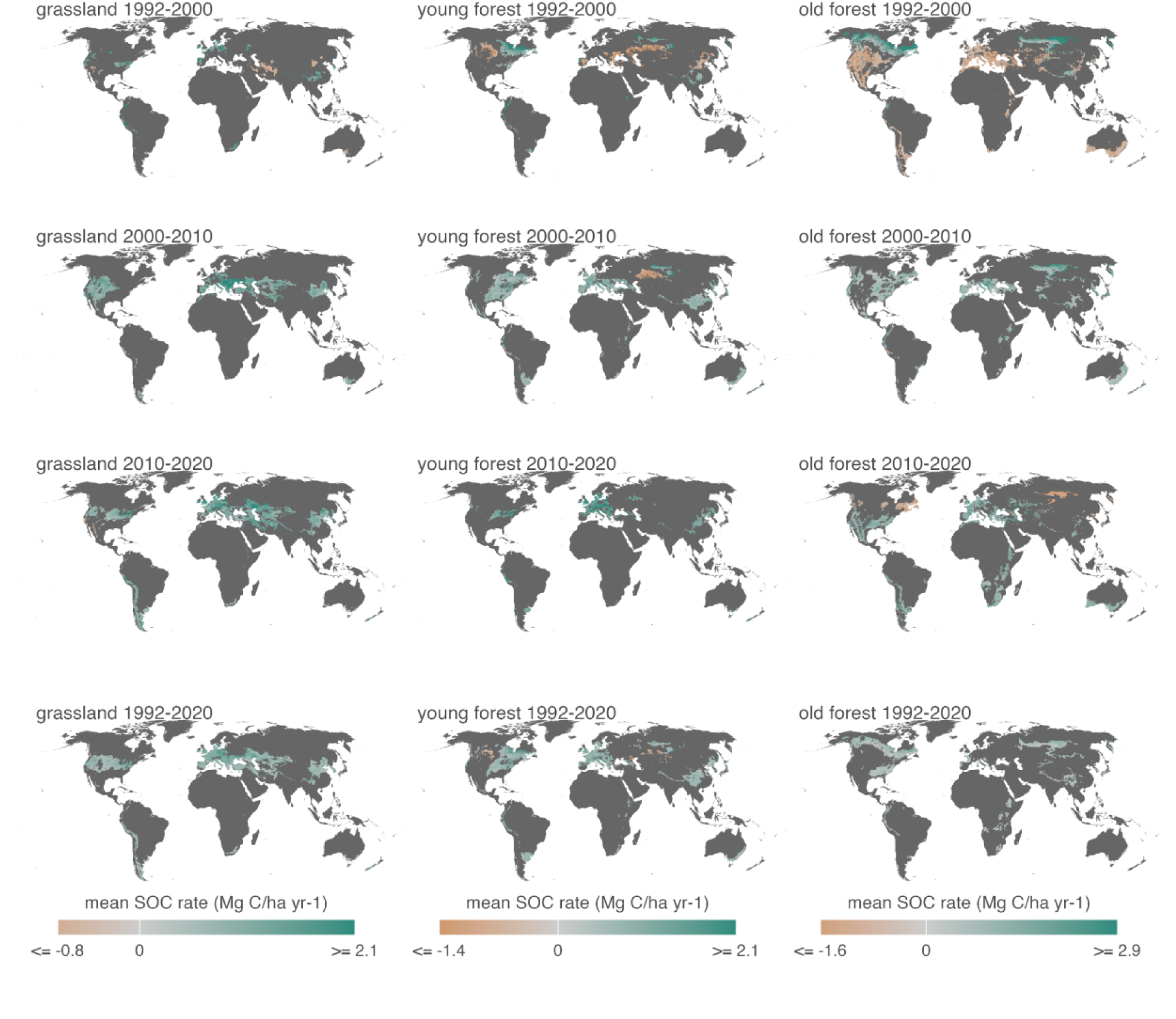
Maps of predicted pixel-level SOC rates (posterior means), showing only the pixels where SOC increased or decreased with more than 97.5% posterior probability.

**Extended Data Fig. 6.**
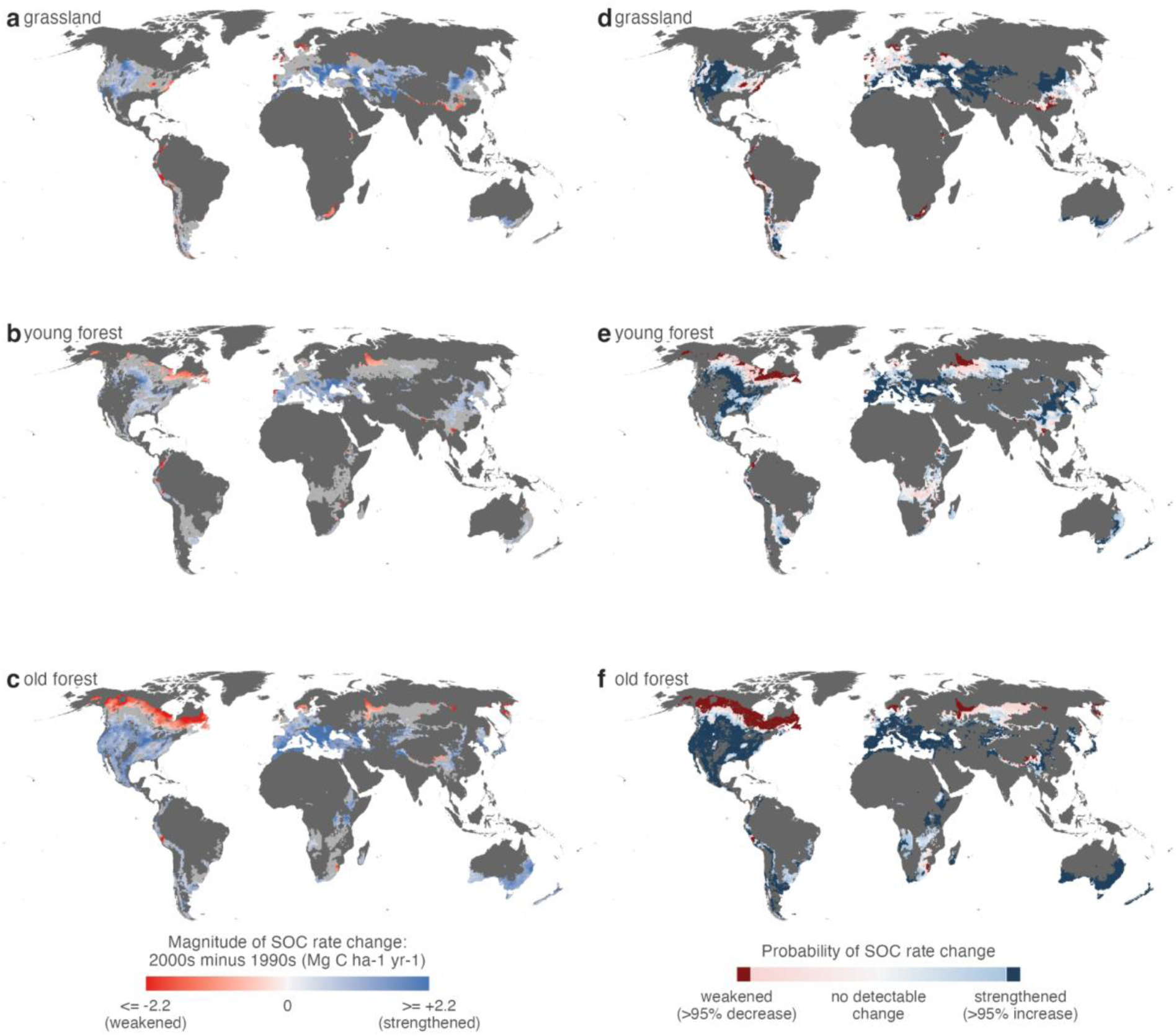
Strengthening or weakening of the SOC sink at pixel level between the 1990s and 2000s. Panels a–c show the magnitude of posterior-mean SOC rate change between the two decades, visualizing only pixels with detectable strengthening or weakening. Other in-domain pixels are shown as light gray. The SOC sink strengthened (blue) for a pixel if the SOC rate increased from the previous decade, and weakened (red) if it decreased. Panels d–f show the probability of SOC rate change. Strengthening or weakening is detectable if more than 95% of the posterior predictions show the same direction of SOC rate change (darkened in panels d–f).

**Extended Data Fig. 7.**
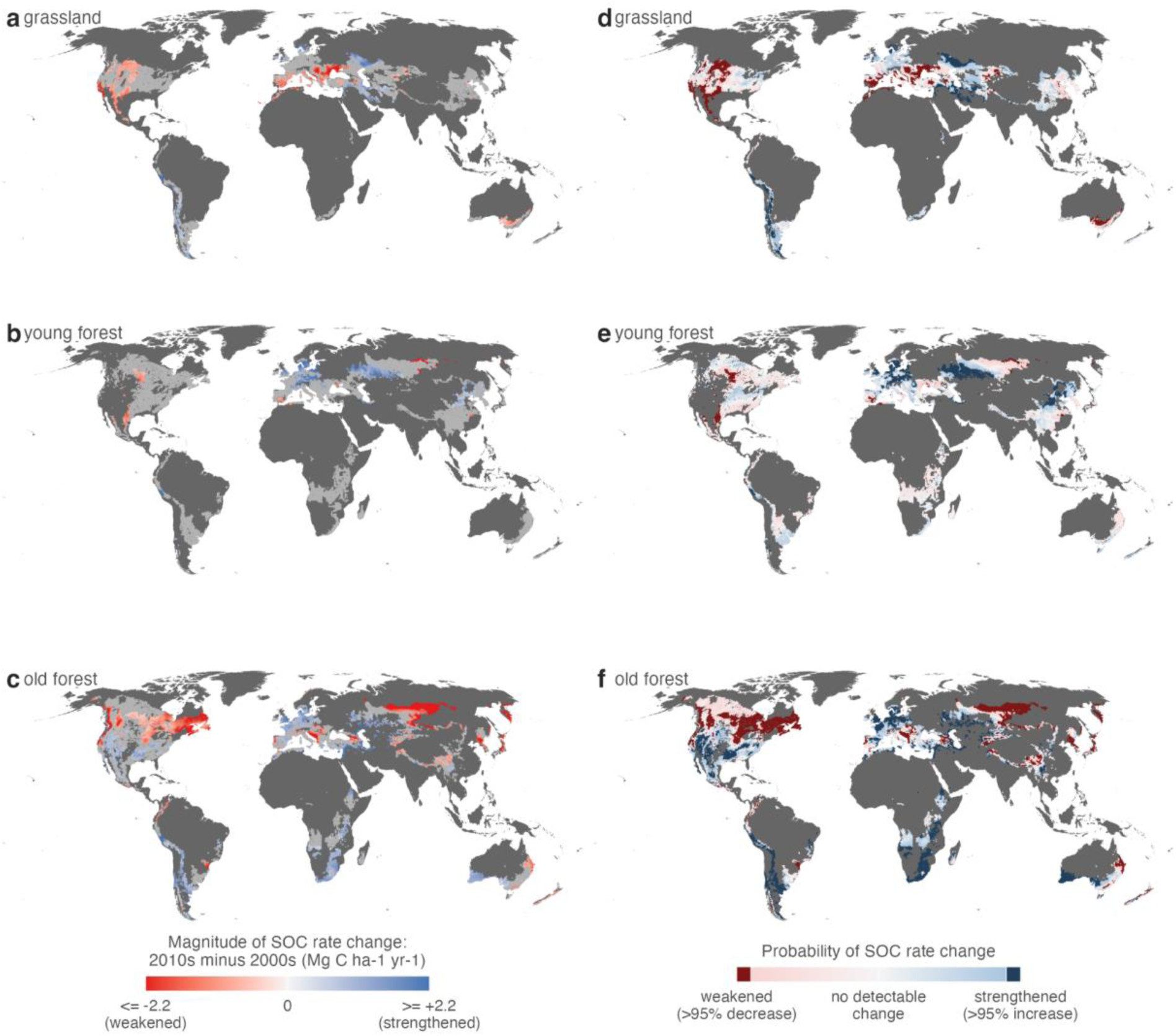
Strengthening or weakening of the SOC sink at pixel level between the 2000s and 2010s. Panels a–c show the magnitude of posterior-mean SOC rate change between the two decades, visualizing only pixels with detectable strengthening or weakening. Other in-domain pixels are shown as light gray. The SOC sink strengthened (blue) for a pixel if the SOC rate increased from the previous decade, and weakened (red) if it decreased. Panels d–f show the probability of SOC rate change. Strengthening or weakening is detectable if more than 95% of the posterior predictions show the same direction of SOC rate change (darkened in panels d–f).

**Extended Data Table 1.**
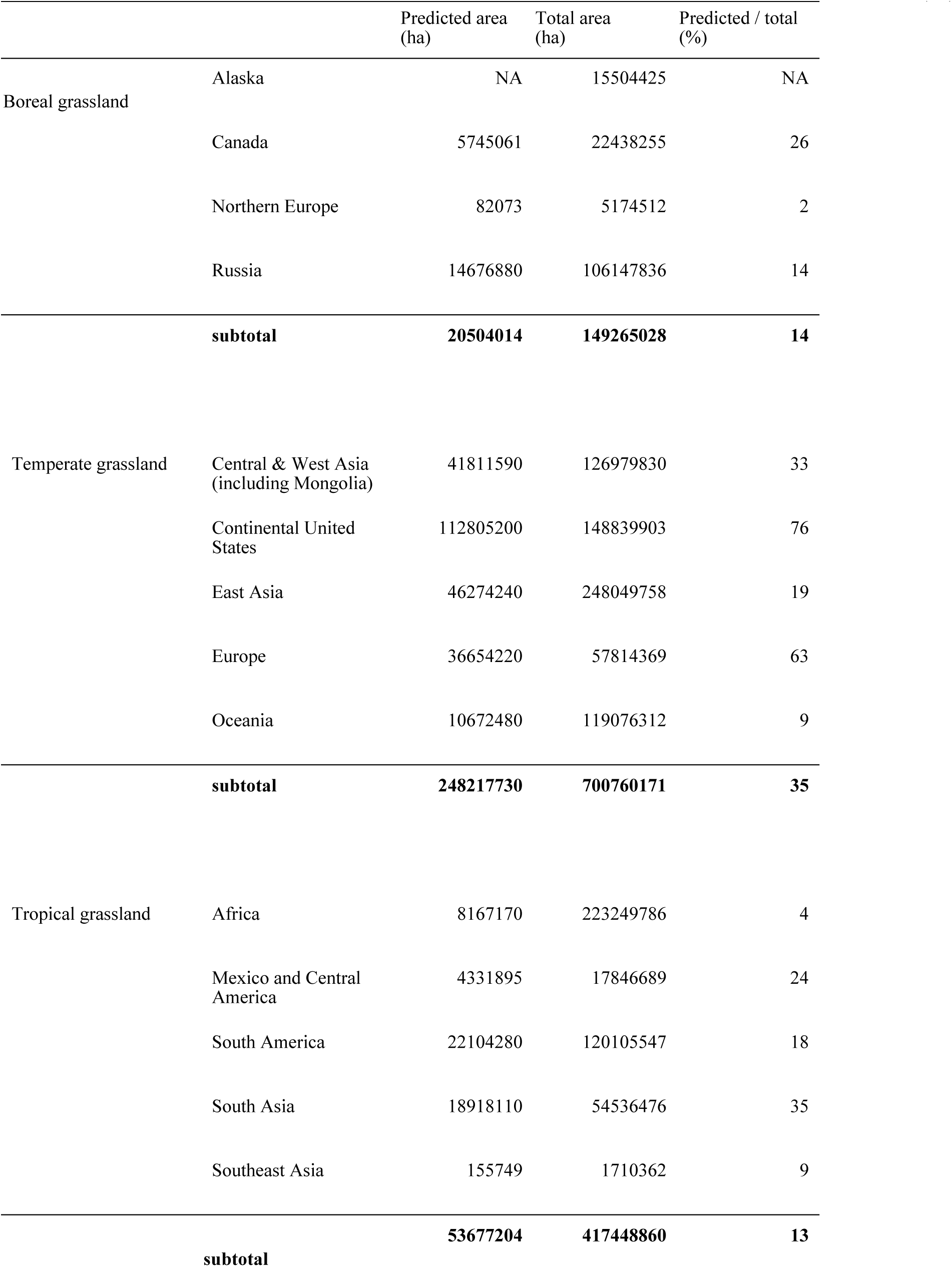
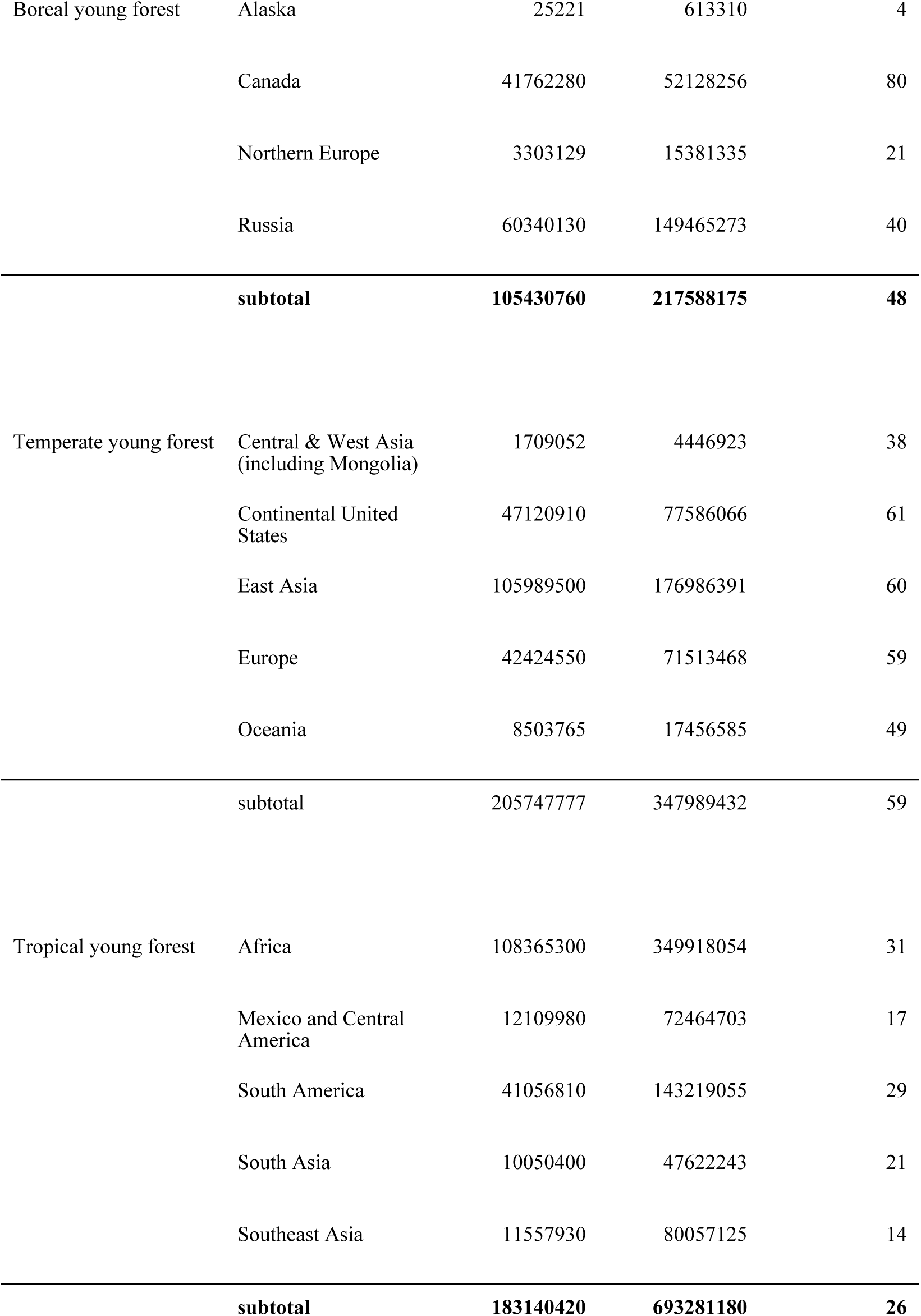
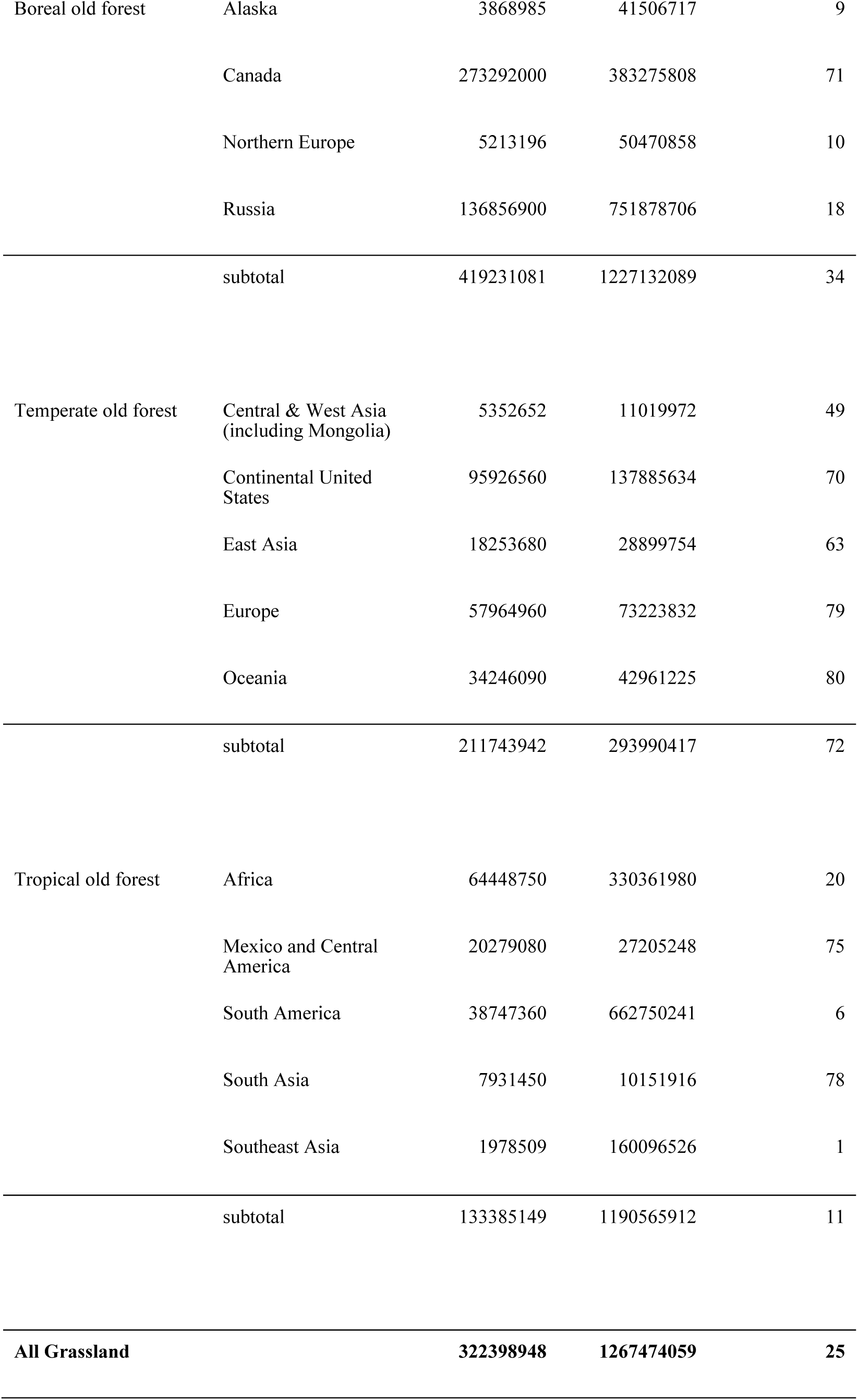
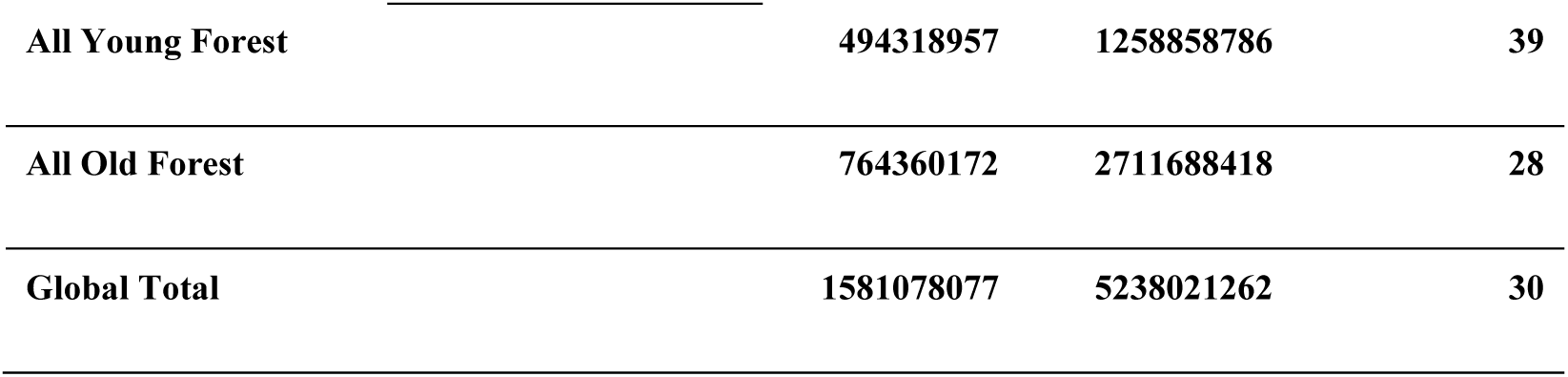
Prediction (in-domain) area applied to each land cover (ha), total area without land cover change 1992–2020 in these regions (ha), and the percentage of predicted area in all land cover area (%).

|  |  | Predicted area (ha) | Total area (ha) | Predicted / total (%) |
|---|---|---|---|---|
| Boreal grassland | Alaska | NA | 15504425 | NA |
|  | Canada | 5745061 | 22438255 | 26 |
|  | Northern Europe | 82073 | 5174512 | 2 |
|  | Russia | 14676880 | 106147836 | 14 |
|  | **subtotal** | **20504014** | **149265028** | **14** |
| Temperate grassland | Central & West Asia (including Mongolia) | 41811590 | 126979830 | 33 |
|  | Continental United States | 112805200 | 148839903 | 76 |
|  | East Asia | 46274240 | 248049758 | 19 |
|  | Europe | 36654220 | 57814369 | 63 |
|  | Oceania | 10672480 | 119076312 | 9 |
|  | **subtotal** | **248217730** | **700760171** | **35** |
| Tropical grassland | Africa | 8167170 | 223249786 | 4 |
|  | Mexico and Central America | 4331895 | 17846689 | 24 |
|  | South America | 22104280 | 120105547 | 18 |
|  | South Asia | 18918110 | 54536476 | 35 |
|  | Southeast Asia | 155749 | 1710362 | 9 |
|  | **subtotal** | **53677204** | **417448860** | **13** |
| Boreal young forest | Alaska | 25221 | 613310 | 4 |
|  | Canada | 41762280 | 52128256 | 80 |
|  | Northern Europe | 3303129 | 15381335 | 21 |
|  | Russia | 60340130 | 149465273 | 40 |
|  | **subtotal** | **105430760** | **217588175** | **48** |
| Temperate young forest | Central & West Asia (including Mongolia) | 1709052 | 4446923 | 38 |
|  | Continental United States | 47120910 | 77586066 | 61 |
|  | East Asia | 105989500 | 176986391 | 60 |
|  | Europe | 42424550 | 71513468 | 59 |
|  | Oceania | 8503765 | 17456585 | 49 |
|  | subtotal | 205747777 | 347989432 | 59 |
| Tropical young forest | Africa | 108365300 | 349918054 | 31 |
|  | Mexico and Central America | 12109980 | 72464703 | 17 |
|  | South America | 41056810 | 143219055 | 29 |
|  | South Asia | 10050400 | 47622243 | 21 |
|  | Southeast Asia | 11557930 | 80057125 | 14 |
|  | **subtotal** | **183140420** | **693281180** | **26** |
| Boreal old forest | Alaska | 3868985 | 41506717 | 9 |
|  | Canada | 273292000 | 383275808 | 71 |
|  | Northern Europe | 5213196 | 50470858 | 10 |
|  | Russia | 136856900 | 751878706 | 18 |
|  | subtotal | 419231081 | 1227132089 | 34 |
| Temperate old forest | Central & West Asia (including Mongolia) | 5352652 | 11019972 | 49 |
|  | Continental United States | 95926560 | 137885634 | 70 |
|  | East Asia | 18253680 | 28899754 | 63 |
|  | Europe | 57964960 | 73223832 | 79 |
|  | Oceania | 34246090 | 42961225 | 80 |
|  | subtotal | 211743942 | 293990417 | 72 |
| Tropical old forest | Africa | 64448750 | 330361980 | 20 |
|  | Mexico and Central America | 20279080 | 27205248 | 75 |
|  | South America | 38747360 | 662750241 | 6 |
|  | South Asia | 7931450 | 10151916 | 78 |
|  | Southeast Asia | 1978509 | 160096526 | 1 |
|  | subtotal | 133385149 | 1190565912 | 11 |
| **All Grassland** |  | **322398948** | **1267474059** | **25** |
| **All Young Forest** |  | **494318957** | **1258858786** | **39** |
| **All Old Forest** |  | **764360172** | **2711688418** | **28** |
| **Global Total** |  | **1581078077** | **5238021262** | **30** |

**Extended Data Table 2.**
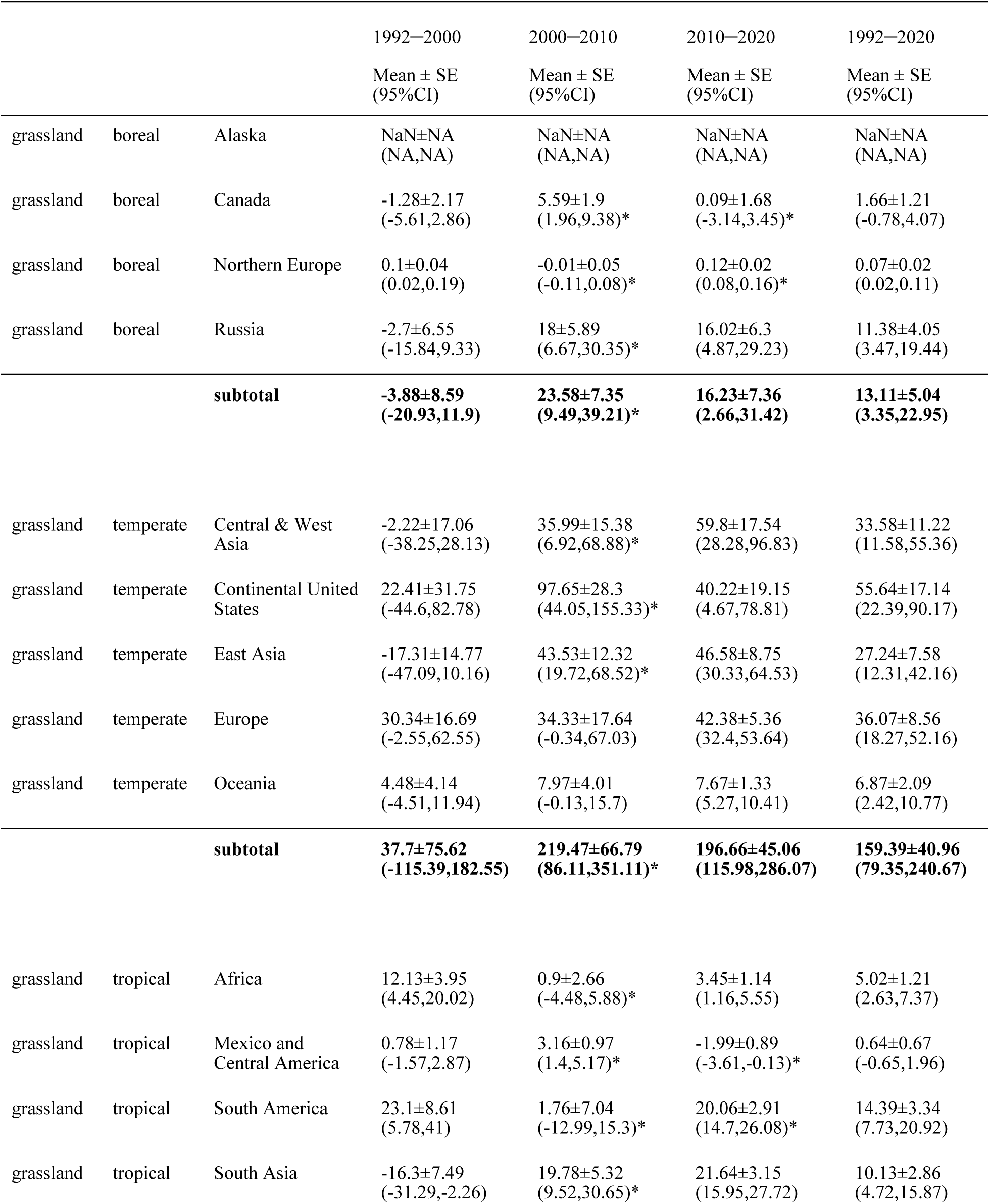
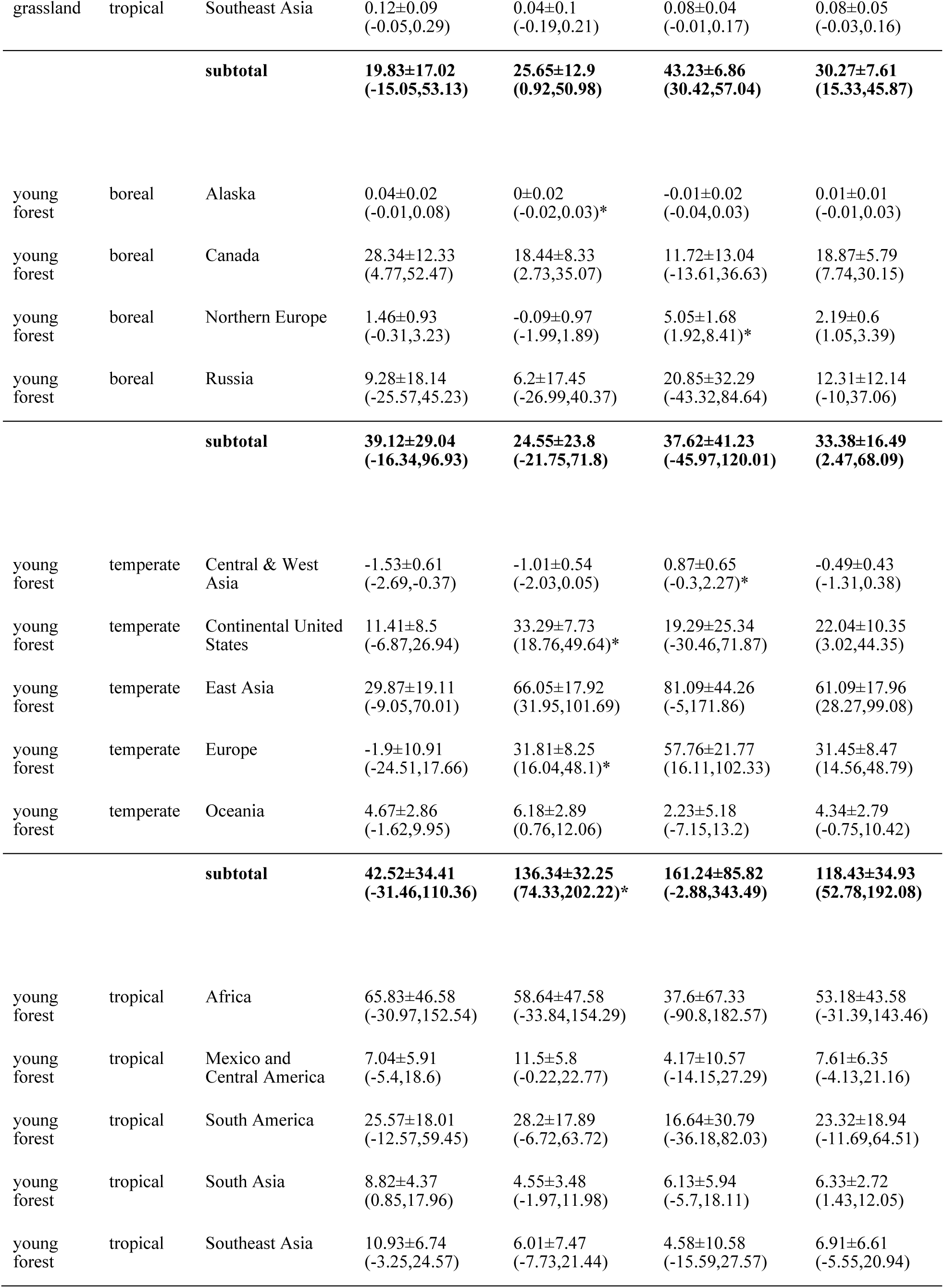
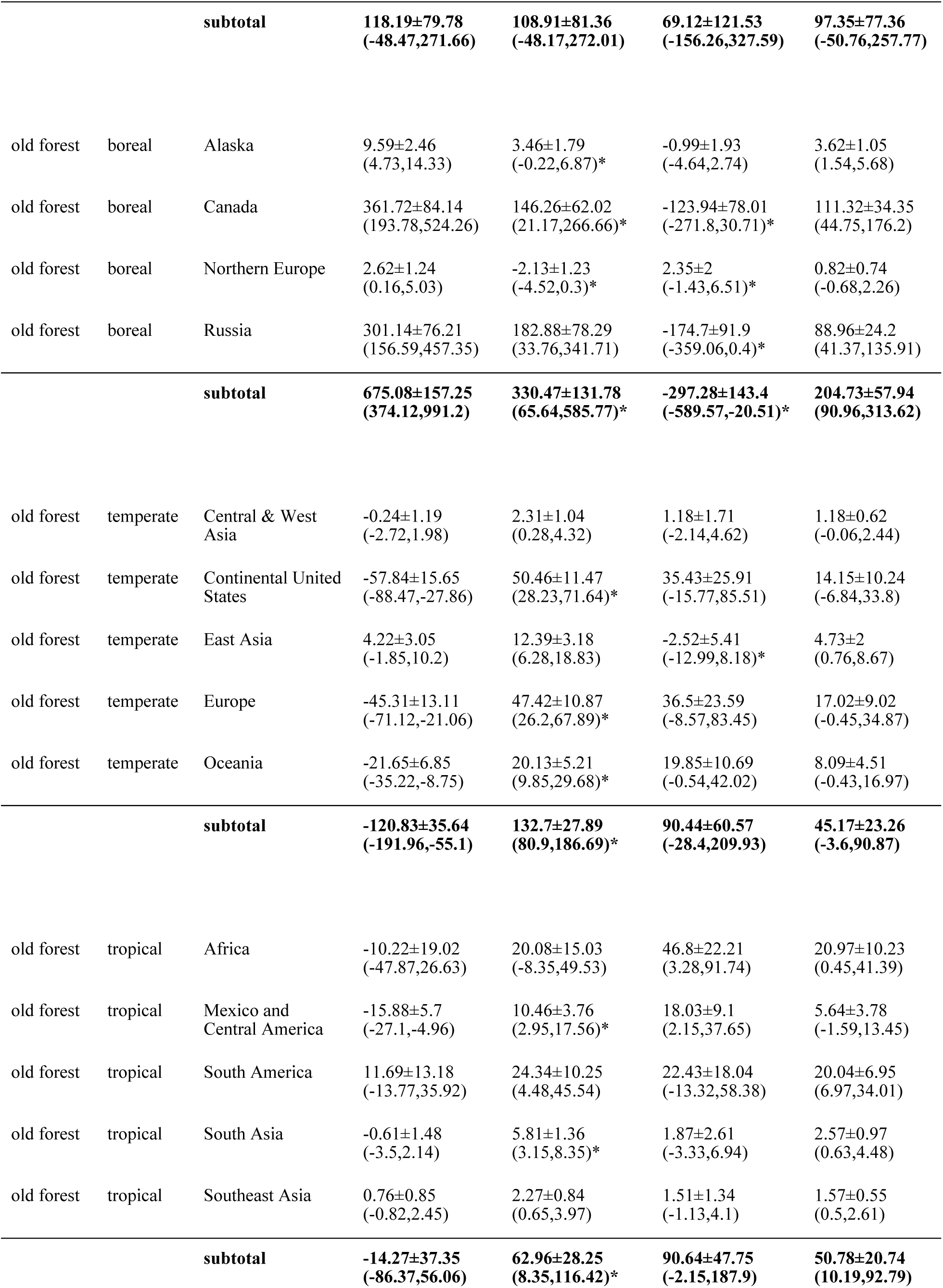
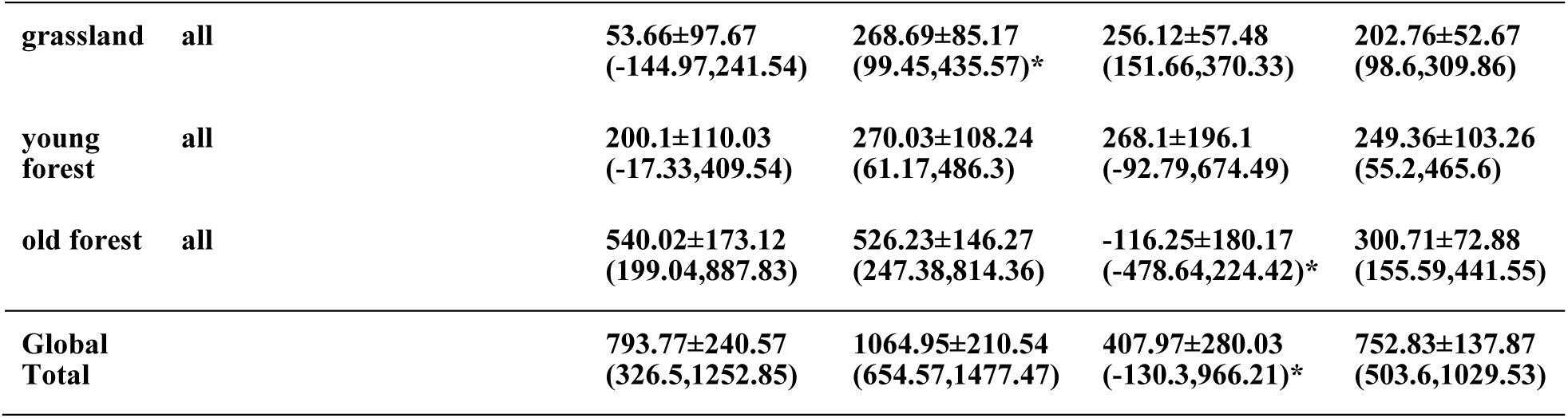
SOC stock changes by region (Unit: Tg C yr^-1^). Asterisks in 2000–2010, 2010–2020 columns indicate the difference in total SOC change from the previous decade is detectable (>95% posterior probability of strengthening or weakening).

|  |  |  | 1992─2000 | 2000─2010 | 2010─2020 | 1992─2020 |
|---|---|---|---|---|---|---|
|  |  |  | Mean ± SE<br>(95%CI) | Mean ± SE<br>(95%CI) | Mean ± SE<br>(95%CI) | Mean ± SE<br>(95%CI) |
| grassland | boreal | Alaska | NaN±NA<br>(NA,NA) | NaN±NA<br>(NA,NA) | NaN±NA<br>(NA,NA) | NaN±NA<br>(NA,NA) |
| grassland | boreal | Canada | -1.28±2.17<br>(-5.61,2.86) | 5.59±1.9<br>(1.96,9.38)* | 0.09±1.68<br>(-3.14,3.45)* | 1.66±1.21<br>(-0.78,4.07) |
| grassland | boreal | Northern Europe | 0.1±0.04<br>(0.02,0.19) | -0.01±0.05<br>(-0.11,0.08)* | 0.12±0.02<br>(0.08,0.16)* | 0.07±0.02<br>(0.02,0.11) |
| grassland | boreal | Russia | -2.7±6.55<br>(-15.84,9.33) | 18±5.89<br>(6.67,30.35)* | 16.02±6.3<br>(4.87,29.23) | 11.38±4.05<br>(3.47,19.44) |
|  |  | **subtotal** | **-3.88±8.59<br>(-20.93,11.9)** | **23.58±7.35<br>(9.49,39.21)\*** | **16.23±7.36<br>(2.66,31.42)** | **13.11±5.04<br>(3.35,22.95)** |
| grassland | temperate | Central & West Asia | -2.22±17.06<br>(-38.25,28.13) | 35.99±15.38<br>(6.92,68.88)* | 59.8±17.54<br>(28.28,96.83) | 33.58±11.22<br>(11.58,55.36) |
| grassland | temperate | Continental United States | 22.41±31.75<br>(-44.6,82.78) | 97.65±28.3<br>(44.05,155.33)* | 40.22±19.15<br>(4.67,78.81) | 55.64±17.14<br>(22.39,90.17) |
| grassland | temperate | East Asia | -17.31±14.77<br>(-47.09,10.16) | 43.53±12.32<br>(19.72,68.52)* | 46.58±8.75<br>(30.33,64.53) | 27.24±7.58<br>(12.31,42.16) |
| grassland | temperate | Europe | 30.34±16.69<br>(-2.55,62.55) | 34.33±17.64<br>(-0.34,67.03) | 42.38±5.36<br>(32.4,53.64) | 36.07±8.56<br>(18.27,52.16) |
| grassland | temperate | Oceania | 4.48±4.14<br>(-4.51,11.94) | 7.97±4.01<br>(-0.13,15.7) | 7.67±1.33<br>(5.27,10.41) | 6.87±2.09<br>(2.42,10.77) |
|  |  | **subtotal** | **37.7±75.62<br>(-115.39,182.55)** | **219.47±66.79<br>(86.11,351.11)\*** | **196.66±45.06<br>(115.98,286.07)** | **159.39±40.96<br>(79.35,240.67)** |
| grassland | tropical | Africa | 12.13±3.95<br>(4.45,20.02) | 0.9±2.66<br>(-4.48,5.88)* | 3.45±1.14<br>(1.16,5.55) | 5.02±1.21<br>(2.63,7.37) |
| grassland | tropical | Mexico and Central America | 0.78±1.17<br>(-1.57,2.87) | 3.16±0.97<br>(1.4,5.17)* | -1.99±0.89<br>(-3.61,-0.13)* | 0.64±0.67<br>(-0.65,1.96) |
| grassland | tropical | South America | 23.1±8.61<br>(5.78,41) | 1.76±7.04<br>(-12.99,15.3)* | 20.06±2.91<br>(14.7,26.08)* | 14.39±3.34<br>(7.73,20.92) |
| grassland | tropical | South Asia | -16.3±7.49<br>(-31.29,-2.26) | 19.78±5.32<br>(9.52,30.65)* | 21.64±3.15<br>(15.95,27.72) | 10.13±2.86<br>(4.72,15.87) |
| grassland | tropical | Southeast Asia | 0.12±0.09<br>(-0.05,0.29) | 0.04±0.1<br>(-0.19,0.21) | 0.08±0.04<br>(-0.01,0.17) | 0.08±0.05<br>(-0.03,0.16) |
|  |  | **subtotal** | **19.83±17.02<br>(-15.05,53.13)** | **25.65±12.9<br>(0.92,50.98)** | **43.23±6.86<br>(30.42,57.04)** | **30.27±7.61<br>(15.33,45.87)** |
| young forest | boreal | Alaska | 0.04±0.02<br>(-0.01,0.08) | 0±0.02<br>(-0.02,0.03)* | -0.01±0.02<br>(-0.04,0.03) | 0.01±0.01<br>(-0.01,0.03) |
| young forest | boreal | Canada | 28.34±12.33<br>(4.77,52.47) | 18.44±8.33<br>(2.73,35.07) | 11.72±13.04<br>(-13.61,36.63) | 18.87±5.79<br>(7.74,30.15) |
| young forest | boreal | Northern Europe | 1.46±0.93<br>(-0.31,3.23) | -0.09±0.97<br>(-1.99,1.89) | 5.05±1.68<br>(1.92,8.41)* | 2.19±0.6<br>(1.05,3.39) |
| young forest | boreal | Russia | 9.28±18.14<br>(-25.57,45.23) | 6.2±17.45<br>(-26.99,40.37) | 20.85±32.29<br>(-43.32,84.64) | 12.31±12.14<br>(-10,37.06) |
|  |  | **subtotal** | **39.12±29.04<br>(-16.34,96.93)** | **24.55±23.8<br>(-21.75,71.8)** | **37.62±41.23<br>(-45.97,120.01)** | **33.38±16.49<br>(2.47,68.09)** |
| young forest | temperate | Central & West Asia | -1.53±0.61<br>(-2.69,-0.37) | -1.01±0.54<br>(-2.03,0.05) | 0.87±0.65<br>(-0.3,2.27)* | -0.49±0.43<br>(-1.31,0.38) |
| young forest | temperate | Continental United States | 11.41±8.5<br>(-6.87,26.94) | 33.29±7.73<br>(18.76,49.64)* | 19.29±25.34<br>(-30.46,71.87) | 22.04±10.35<br>(3.02,44.35) |
| young forest | temperate | East Asia | 29.87±19.11<br>(-9.05,70.01) | 66.05±17.92<br>(31.95,101.69) | 81.09±44.26<br>(-5,171.86) | 61.09±17.96<br>(28.27,99.08) |
| young forest | temperate | Europe | -1.9±10.91<br>(-24.51,17.66) | 31.81±8.25<br>(16.04,48.1)* | 57.76±21.77<br>(16.11,102.33) | 31.45±8.47<br>(14.56,48.79) |
| young forest | temperate | Oceania | 4.67±2.86<br>(-1.62,9.95) | 6.18±2.89<br>(0.76,12.06) | 2.23±5.18<br>(-7.15,13.2) | 4.34±2.79<br>(-0.75,10.42) |
|  |  | **subtotal** | **42.52±34.41<br>(-31.46,110.36)** | **136.34±32.25<br>(74.33,202.22)\*** | **161.24±85.82<br>(-2.88,343.49)** | **118.43±34.93<br>(52.78,192.08)** |
| young forest | tropical | Africa | 65.83±46.58<br>(-30.97,152.54) | 58.64±47.58<br>(-33.84,154.29) | 37.6±67.33<br>(-90.8,182.57) | 53.18±43.58<br>(-31.39,143.46) |
| young forest | tropical | Mexico and Central America | 7.04±5.91<br>(-5.4,18.6) | 11.5±5.8<br>(-0.22,22.77) | 4.17±10.57<br>(-14.15,27.29) | 7.61±6.35<br>(-4.13,21.16) |
| young forest | tropical | South America | 25.57±18.01<br>(-12.57,59.45) | 28.2±17.89<br>(-6.72,63.72) | 16.64±30.79<br>(-36.18,82.03) | 23.32±18.94<br>(-11.69,64.51) |
| young forest | tropical | South Asia | 8.82±4.37<br>(0.85,17.96) | 4.55±3.48<br>(-1.97,11.98) | 6.13±5.94<br>(-5.7,18.11) | 6.33±2.72<br>(1.43,12.05) |
| young forest | tropical | Southeast Asia | 10.93±6.74<br>(-3.25,24.57) | 6.01±7.47<br>(-7.73,21.44) | 4.58±10.58<br>(-15.59,27.57) | 6.91±6.61<br>(-5.55,20.94) |
|  |  | **subtotal** | **118.19±79.78<br>(-48.47,271.66)** | **108.91±81.36<br>(-48.17,272.01)** | **69.12±121.53<br>(-156.26,327.59)** | **97.35±77.36<br>(-50.76,257.77)** |
| old forest | boreal | Alaska | 9.59±2.46<br>(4.73,14.33) | 3.46±1.79<br>(-0.22,6.87)* | -0.99±1.93<br>(-4.64,2.74) | 3.62±1.05<br>(1.54,5.68) |
| old forest | boreal | Canada | 361.72±84.14<br>(193.78,524.26) | 146.26±62.02<br>(21.17,266.66)* | -123.94±78.01<br>(-271.8,30.71)* | 111.32±34.35<br>(44.75,176.2) |
| old forest | boreal | Northern Europe | 2.62±1.24<br>(0.16,5.03) | -2.13±1.23<br>(-4.52,0.3)* | 2.35±2<br>(-1.43,6.51)* | 0.82±0.74<br>(-0.68,2.26) |
| old forest | boreal | Russia | 301.14±76.21<br>(156.59,457.35) | 182.88±78.29<br>(33.76,341.71) | -174.7±91.9<br>(-359.06,0.4)* | 88.96±24.2<br>(41.37,135.91) |
|  |  | **subtotal** | **675.08±157.25<br>(374.12,991.2)** | **330.47±131.78<br>(65.64,585.77)\*** | **-297.28±143.4<br>(-589.57,-20.51)\*** | **204.73±57.94<br>(90.96,313.62)** |
| old forest | temperate | Central & West Asia | -0.24±1.19<br>(-2.72,1.98) | 2.31±1.04<br>(0.28,4.32) | 1.18±1.71<br>(-2.14,4.62) | 1.18±0.62<br>(-0.06,2.44) |
| old forest | temperate | Continental United States | -57.84±15.65<br>(-88.47,-27.86) | 50.46±11.47<br>(28.23,71.64)* | 35.43±25.91<br>(-15.77,85.51) | 14.15±10.24<br>(-6.84,33.8) |
| old forest | temperate | East Asia | 4.22±3.05<br>(-1.85,10.2) | 12.39±3.18<br>(6.28,18.83) | -2.52±5.41<br>(-12.99,8.18)* | 4.73±2<br>(0.76,8.67) |
| old forest | temperate | Europe | -45.31±13.11<br>(-71.12,-21.06) | 47.42±10.87<br>(26.2,67.89)* | 36.5±23.59<br>(-8.57,83.45) | 17.02±9.02<br>(-0.45,34.87) |
| old forest | temperate | Oceania | -21.65±6.85<br>(-35.22,-8.75) | 20.13±5.21<br>(9.85,29.68)* | 19.85±10.69<br>(-0.54,42.02) | 8.09±4.51<br>(-0.43,16.97) |
|  |  | **subtotal** | **-120.83±35.64<br>(-191.96,-55.1)** | **132.7±27.89<br>(80.9,186.69)\*** | **90.44±60.57<br>(-28.4,209.93)** | **45.17±23.26<br>(-3.6,90.87)** |
| old forest | tropical | Africa | -10.22±19.02<br>(-47.87,26.63) | 20.08±15.03<br>(-8.35,49.53) | 46.8±22.21<br>(3.28,91.74) | 20.97±10.23<br>(0.45,41.39) |
| old forest | tropical | Mexico and Central America | -15.88±5.7<br>(-27.1,-4.96) | 10.46±3.76<br>(2.95,17.56)* | 18.03±9.1<br>(2.15,37.65) | 5.64±3.78<br>(-1.59,13.45) |
| old forest | tropical | South America | 11.69±13.18<br>(-13.77,35.92) | 24.34±10.25<br>(4.48,45.54) | 22.43±18.04<br>(-13.32,58.38) | 20.04±6.95<br>(6.97,34.01) |
| old forest | tropical | South Asia | -0.61±1.48<br>(-3.5,2.14) | 5.81±1.36<br>(3.15,8.35)* | 1.87±2.61<br>(-3.33,6.94) | 2.57±0.97<br>(0.63,4.48) |
| old forest | tropical | Southeast Asia | 0.76±0.85<br>(-0.82,2.45) | 2.27±0.84<br>(0.65,3.97) | 1.51±1.34<br>(-1.13,4.1) | 1.57±0.55<br>(0.5,2.61) |
|  |  | **subtotal** | **-14.27±37.35<br>(-86.37,56.06)** | **62.96±28.25<br>(8.35,116.42)\*** | **90.64±47.75<br>(-2.15,187.9)** | **50.78±20.74<br>(10.19,92.79)** |
| **grassland** | **all** |  | **53.66±97.67<br>(-144.97,241.54)** | **268.69±85.17<br>(99.45,435.57)\*** | **256.12±57.48<br>(151.66,370.33)** | **202.76±52.67<br>(98.6,309.86)** |
| **young forest** | **all** |  | **200.1±110.03<br>(-17.33,409.54)** | **270.03±108.24<br>(61.17,486.3)** | **268.1±196.1<br>(-92.79,674.49)** | **249.36±103.26<br>(55.2,465.6)** |
| **old forest** | **all** |  | **540.02±173.12<br>(199.04,887.83)** | **526.23±146.27<br>(247.38,814.36)** | **-116.25±180.17<br>(-478.64,224.42)\*** | **300.71±72.88<br>(155.59,441.55)** |
| **Global Total** |  |  | **793.77±240.57<br>(326.5,1252.85)** | **1064.95±210.54<br>(654.57,1477.47)** | **407.97±280.03<br>(-130.3,966.21)\*** | **752.83±137.87<br>(503.6,1029.53)** |

**Extended Data Table 3.**
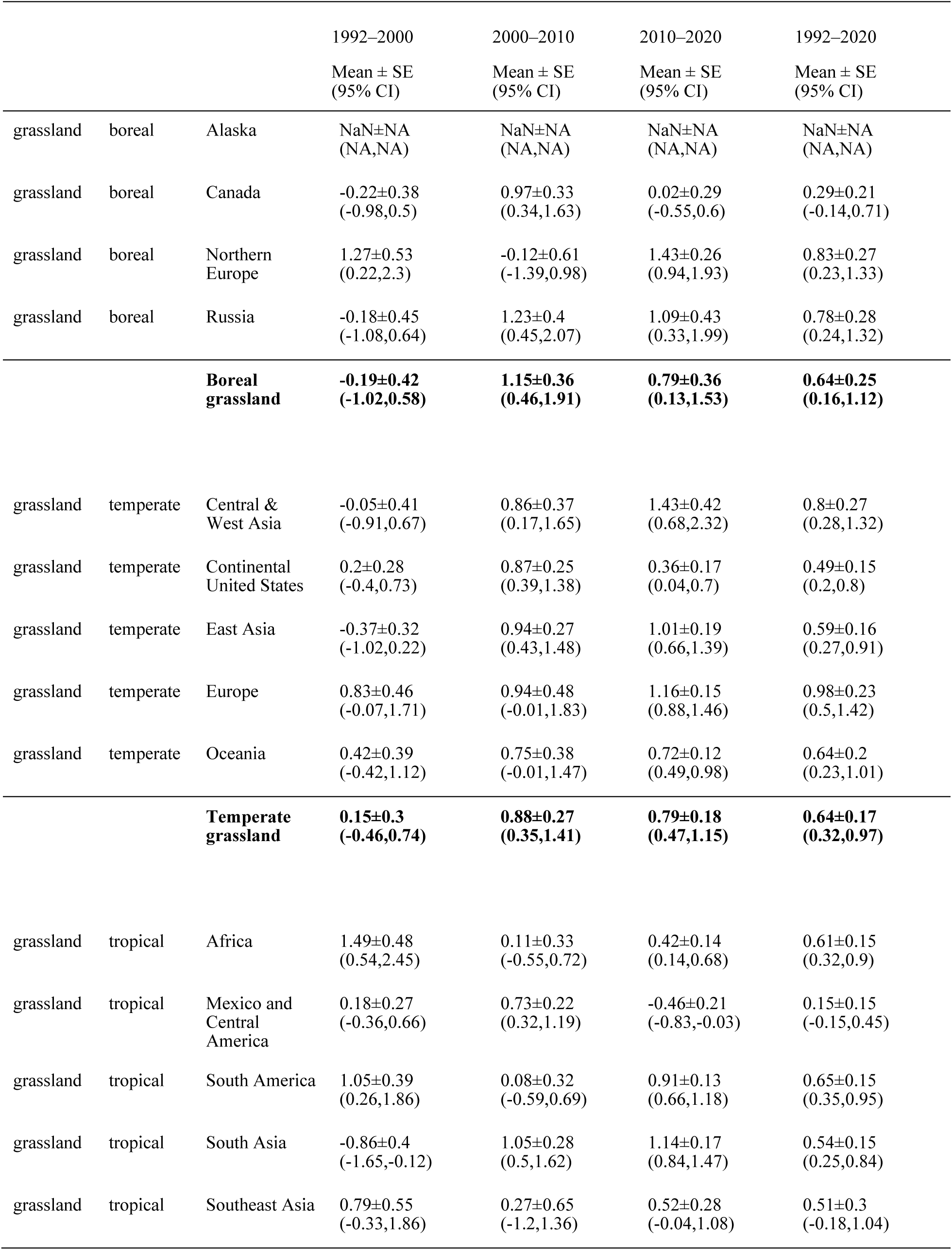
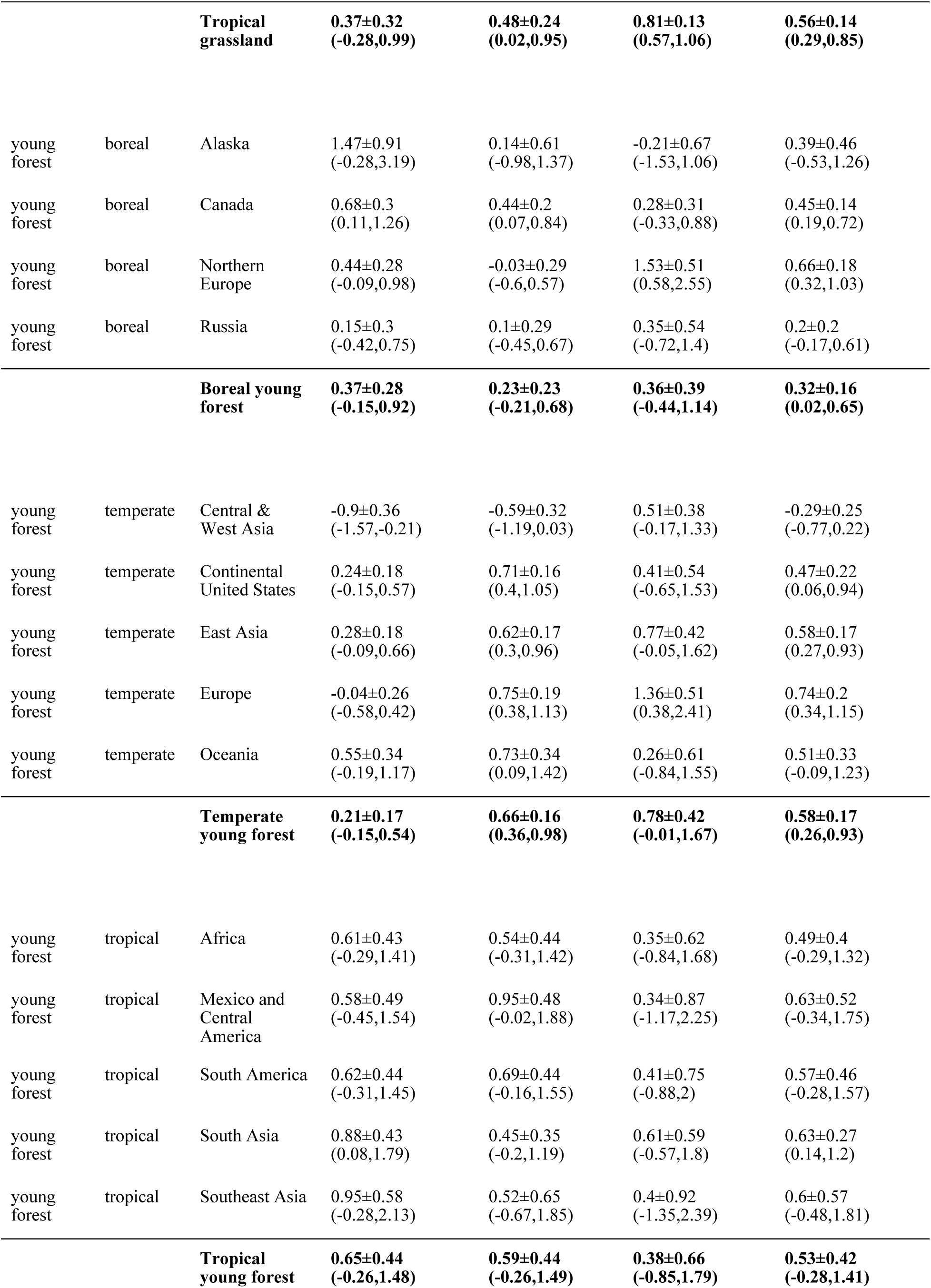
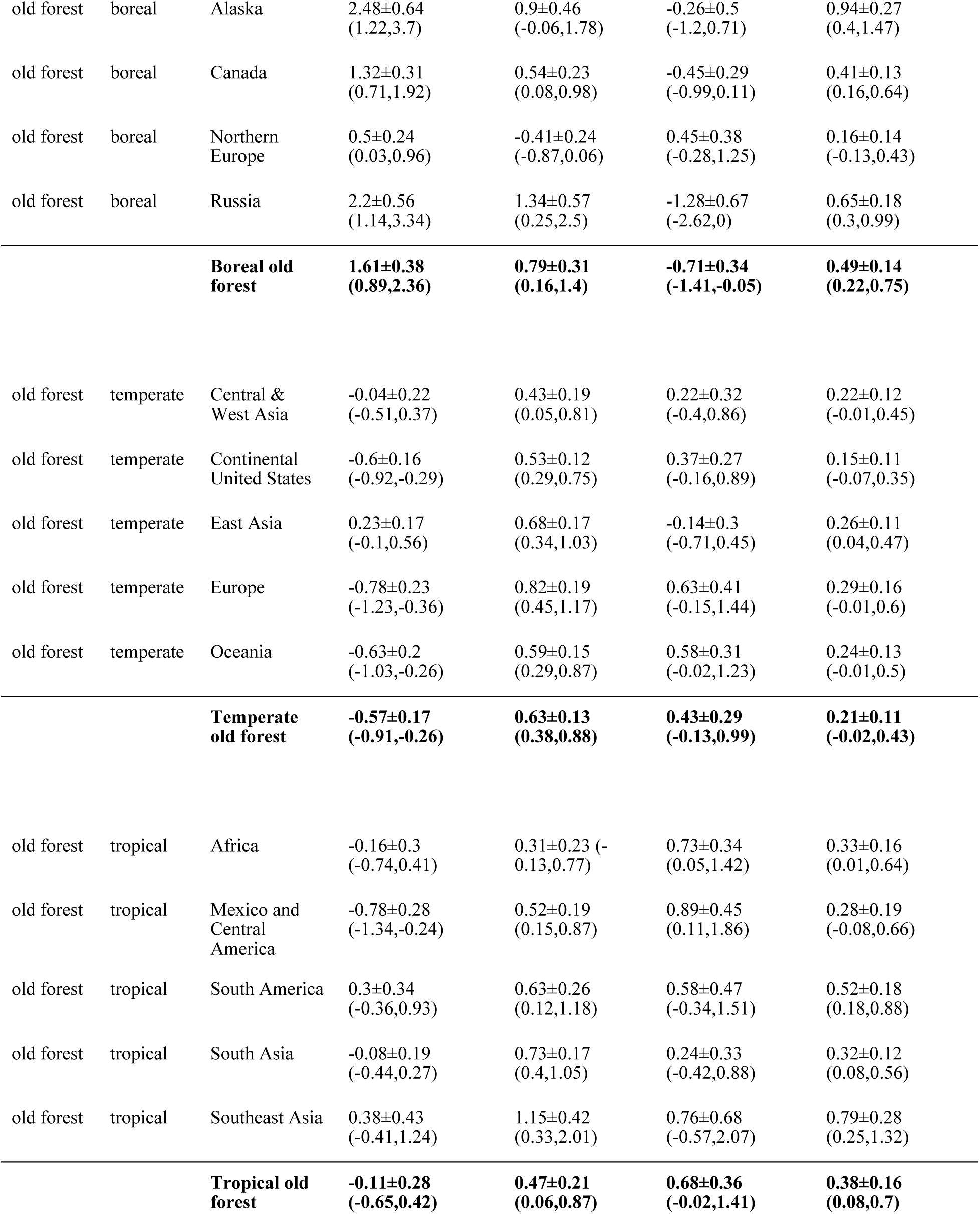
Average SOC change per area by region and biome (Unit: Mg C ha^-1^ yr^-1^).

|  |  |  | 1992–2000 | 2000–2010 | 2010–2020 | 1992–2020 |
|---|---|---|---|---|---|---|
|  |  |  | Mean ± SE (95% CI) | Mean ± SE (95% CI) | Mean ± SE (95% CI) | Mean ± SE (95% CI) |
| grassland | boreal | Alaska | NaN±NA (NA,NA) | NaN±NA (NA,NA) | NaN±NA (NA,NA) | NaN±NA (NA,NA) |
| grassland | boreal | Canada | -0.22±0.38 (-0.98,0.5) | 0.97±0.33 (0.34,1.63) | 0.02±0.29 (-0.55,0.6) | 0.29±0.21 (-0.14,0.71) |
| grassland | boreal | Northern Europe | 1.27±0.53 (0.22,2.3) | -0.12±0.61 (-1.39,0.98) | 1.43±0.26 (0.94,1.93) | 0.83±0.27 (0.23,1.33) |
| grassland | boreal | Russia | -0.18±0.45 (-1.08,0.64) | 1.23±0.4 (0.45,2.07) | 1.09±0.43 (0.33,1.99) | 0.78±0.28 (0.24,1.32) |
|  |  | **Boreal grassland** | **-0.19±0.42 (-1.02,0.58)** | **1.15±0.36 (0.46,1.91)** | **0.79±0.36 (0.13,1.53)** | **0.64±0.25 (0.16,1.12)** |
| grassland | temperate | Central & West Asia | -0.05±0.41 (-0.91,0.67) | 0.86±0.37 (0.17,1.65) | 1.43±0.42 (0.68,2.32) | 0.8±0.27 (0.28,1.32) |
| grassland | temperate | Continental United States | 0.2±0.28 (-0.4,0.73) | 0.87±0.25 (0.39,1.38) | 0.36±0.17 (0.04,0.7) | 0.49±0.15 (0.2,0.8) |
| grassland | temperate | East Asia | -0.37±0.32 (-1.02,0.22) | 0.94±0.27 (0.43,1.48) | 1.01±0.19 (0.66,1.39) | 0.59±0.16 (0.27,0.91) |
| grassland | temperate | Europe | 0.83±0.46 (-0.07,1.71) | 0.94±0.48 (-0.01,1.83) | 1.16±0.15 (0.88,1.46) | 0.98±0.23 (0.5,1.42) |
| grassland | temperate | Oceania | 0.42±0.39 (-0.42,1.12) | 0.75±0.38 (-0.01,1.47) | 0.72±0.12 (0.49,0.98) | 0.64±0.2 (0.23,1.01) |
|  |  | **Temperate grassland** | **0.15±0.3 (-0.46,0.74)** | **0.88±0.27 (0.35,1.41)** | **0.79±0.18 (0.47,1.15)** | **0.64±0.17 (0.32,0.97)** |
| grassland | tropical | Africa | 1.49±0.48 (0.54,2.45) | 0.11±0.33 (-0.55,0.72) | 0.42±0.14 (0.14,0.68) | 0.61±0.15 (0.32,0.9) |
| grassland | tropical | Mexico and Central America | 0.18±0.27 (-0.36,0.66) | 0.73±0.22 (0.32,1.19) | -0.46±0.21 (-0.83,-0.03) | 0.15±0.15 (-0.15,0.45) |
| grassland | tropical | South America | 1.05±0.39 (0.26,1.86) | 0.08±0.32 (-0.59,0.69) | 0.91±0.13 (0.66,1.18) | 0.65±0.15 (0.35,0.95) |
| grassland | tropical | South Asia | -0.86±0.4 (-1.65,-0.12) | 1.05±0.28 (0.5,1.62) | 1.14±0.17 (0.84,1.47) | 0.54±0.15 (0.25,0.84) |
| grassland | tropical | Southeast Asia | 0.79±0.55 (-0.33,1.86) | 0.27±0.65 (-1.2,1.36) | 0.52±0.28 (-0.04,1.08) | 0.51±0.3 (-0.18,1.04) |
|  |  | **Tropical grassland** | **0.37±0.32 (-0.28,0.99)** | **0.48±0.24 (0.02,0.95)** | **0.81±0.13 (0.57,1.06)** | **0.56±0.14 (0.29,0.85)** |
| young forest | boreal | Alaska | 1.47±0.91 (-0.28,3.19) | 0.14±0.61 (-0.98,1.37) | -0.21±0.67 (-1.53,1.06) | 0.39±0.46 (-0.53,1.26) |
| young forest | boreal | Canada | 0.68±0.3 (0.11,1.26) | 0.44±0.2 (0.07,0.84) | 0.28±0.31 (-0.33,0.88) | 0.45±0.14 (0.19,0.72) |
| young forest | boreal | Northern Europe | 0.44±0.28 (-0.09,0.98) | -0.03±0.29 (-0.6,0.57) | 1.53±0.51 (0.58,2.55) | 0.66±0.18 (0.32,1.03) |
| young forest | boreal | Russia | 0.15±0.3 (-0.42,0.75) | 0.1±0.29 (-0.45,0.67) | 0.35±0.54 (-0.72,1.4) | 0.2±0.2 (-0.17,0.61) |
|  |  | **Boreal young forest** | **0.37±0.28 (-0.15,0.92)** | **0.23±0.23 (-0.21,0.68)** | **0.36±0.39 (-0.44,1.14)** | **0.32±0.16 (0.02,0.65)** |
| young forest | temperate | Central & West Asia | -0.9±0.36 (-1.57,-0.21) | -0.59±0.32 (-1.19,0.03) | 0.51±0.38 (-0.17,1.33) | -0.29±0.25 (-0.77,0.22) |
| young forest | temperate | Continental United States | 0.24±0.18 (-0.15,0.57) | 0.71±0.16 (0.4,1.05) | 0.41±0.54 (-0.65,1.53) | 0.47±0.22 (0.06,0.94) |
| young forest | temperate | East Asia | 0.28±0.18 (-0.09,0.66) | 0.62±0.17 (0.3,0.96) | 0.77±0.42 (-0.05,1.62) | 0.58±0.17 (0.27,0.93) |
| young forest | temperate | Europe | -0.04±0.26 (-0.58,0.42) | 0.75±0.19 (0.38,1.13) | 1.36±0.51 (0.38,2.41) | 0.74±0.2 (0.34,1.15) |
| young forest | temperate | Oceania | 0.55±0.34 (-0.19,1.17) | 0.73±0.34 (0.09,1.42) | 0.26±0.61 (-0.84,1.55) | 0.51±0.33 (-0.09,1.23) |
|  |  | **Temperate young forest** | **0.21±0.17 (-0.15,0.54)** | **0.66±0.16 (0.36,0.98)** | **0.78±0.42 (-0.01,1.67)** | **0.58±0.17 (0.26,0.93)** |
| young forest | tropical | Africa | 0.61±0.43 (-0.29,1.41) | 0.54±0.44 (-0.31,1.42) | 0.35±0.62 (-0.84,1.68) | 0.49±0.4 (-0.29,1.32) |
| young forest | tropical | Mexico and Central America | 0.58±0.49 (-0.45,1.54) | 0.95±0.48 (-0.02,1.88) | 0.34±0.87 (-1.17,2.25) | 0.63±0.52 (-0.34,1.75) |
| young forest | tropical | South America | 0.62±0.44 (-0.31,1.45) | 0.69±0.44 (-0.16,1.55) | 0.41±0.75 (-0.88,2) | 0.57±0.46 (-0.28,1.57) |
| young forest | tropical | South Asia | 0.88±0.43 (0.08,1.79) | 0.45±0.35 (-0.2,1.19) | 0.61±0.59 (-0.57,1.8) | 0.63±0.27 (0.14,1.2) |
| young forest | tropical | Southeast Asia | 0.95±0.58 (-0.28,2.13) | 0.52±0.65 (-0.67,1.85) | 0.4±0.92 (-1.35,2.39) | 0.6±0.57 (-0.48,1.81) |
|  |  | **Tropical young forest** | **0.65±0.44 (-0.26,1.48)** | **0.59±0.44 (-0.26,1.49)** | **0.38±0.66 (-0.85,1.79)** | **0.53±0.42 (-0.28,1.41)** |
| old forest | boreal | Alaska | 2.48±0.64 (1.22,3.7) | 0.9±0.46 (-0.06,1.78) | -0.26±0.5 (-1.2,0.71) | 0.94±0.27 (0.4,1.47) |
| old forest | boreal | Canada | 1.32±0.31 (0.71,1.92) | 0.54±0.23 (0.08,0.98) | -0.45±0.29 (-0.99,0.11) | 0.41±0.13 (0.16,0.64) |
| old forest | boreal | Northern Europe | 0.5±0.24 (0.03,0.96) | -0.41±0.24 (-0.87,0.06) | 0.45±0.38 (-0.28,1.25) | 0.16±0.14 (-0.13,0.43) |
| old forest | boreal | Russia | 2.2±0.56 (1.14,3.34) | 1.34±0.57 (0.25,2.5) | -1.28±0.67 (-2.62,0) | 0.65±0.18 (0.3,0.99) |
|  |  | **Boreal old forest** | **1.61±0.38 (0.89,2.36)** | **0.79±0.31 (0.16,1.4)** | **-0.71±0.34 (-1.41,-0.05)** | **0.49±0.14 (0.22,0.75)** |
| old forest | temperate | Central & West Asia | -0.04±0.22 (-0.51,0.37) | 0.43±0.19 (0.05,0.81) | 0.22±0.32 (-0.4,0.86) | 0.22±0.12 (-0.01,0.45) |
| old forest | temperate | Continental United States | -0.6±0.16 (-0.92,-0.29) | 0.53±0.12 (0.29,0.75) | 0.37±0.27 (-0.16,0.89) | 0.15±0.11 (-0.07,0.35) |
| old forest | temperate | East Asia | 0.23±0.17 (-0.1,0.56) | 0.68±0.17 (0.34,1.03) | -0.14±0.3 (-0.71,0.45) | 0.26±0.11 (0.04,0.47) |
| old forest | temperate | Europe | -0.78±0.23 (-1.23,-0.36) | 0.82±0.19 (0.45,1.17) | 0.63±0.41 (-0.15,1.44) | 0.29±0.16 (-0.01,0.6) |
| old forest | temperate | Oceania | -0.63±0.2 (-1.03,-0.26) | 0.59±0.15 (0.29,0.87) | 0.58±0.31 (-0.02,1.23) | 0.24±0.13 (-0.01,0.5) |
|  |  | **Temperate old forest** | **-0.57±0.17 (-0.91,-0.26)** | **0.63±0.13 (0.38,0.88)** | **0.43±0.29 (-0.13,0.99)** | **0.21±0.11 (-0.02,0.43)** |
| old forest | tropical | Africa | -0.16±0.3 (-0.74,0.41) | 0.31±0.23 (-0.13,0.77) | 0.73±0.34 (0.05,1.42) | 0.33±0.16 (0.01,0.64) |
| old forest | tropical | Mexico and Central America | -0.78±0.28 (-1.34,-0.24) | 0.52±0.19 (0.15,0.87) | 0.89±0.45 (0.11,1.86) | 0.28±0.19 (-0.08,0.66) |
| old forest | tropical | South America | 0.3±0.34 (-0.36,0.93) | 0.63±0.26 (0.12,1.18) | 0.58±0.47 (-0.34,1.51) | 0.52±0.18 (0.18,0.88) |
| old forest | tropical | South Asia | -0.08±0.19 (-0.44,0.27) | 0.73±0.17 (0.4,1.05) | 0.24±0.33 (-0.42,0.88) | 0.32±0.12 (0.08,0.56) |
| old forest | tropical | Southeast Asia | 0.38±0.43 (-0.41,1.24) | 1.15±0.42 (0.33,2.01) | 0.76±0.68 (-0.57,2.07) | 0.79±0.28 (0.25,1.32) |
|  |  | **Tropical old forest** | **-0.11±0.28 (-0.65,0.42)** | **0.47±0.21 (0.06,0.87)** | **0.68±0.36 (-0.02,1.41)** | **0.38±0.16 (0.08,0.7)** |

## Supplementary Information

**Fig. S1.**
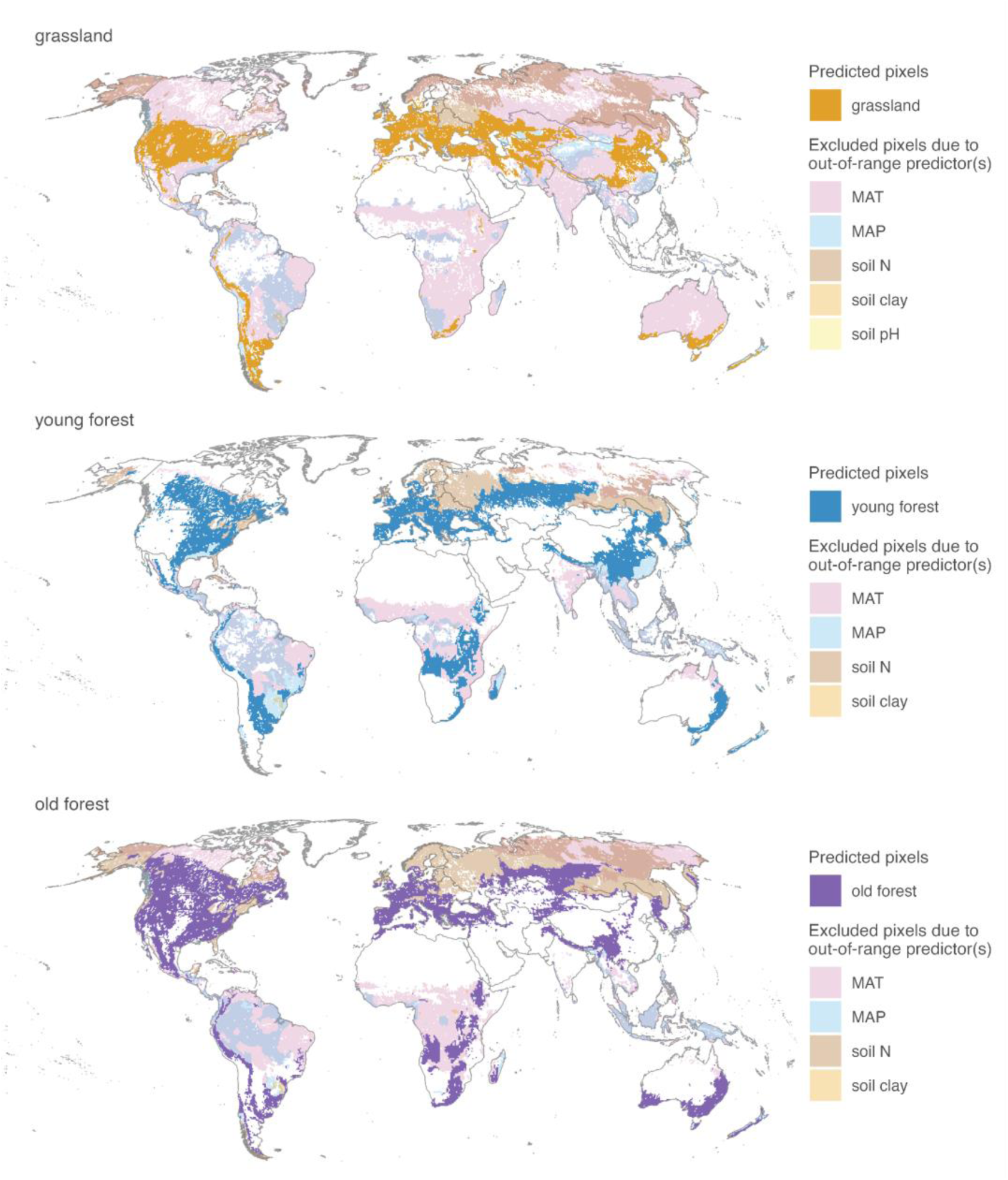
Spatial domain used in upscaling analysis. The pixels within the prediction area should (1) lie within the environmental space supported by observations and (2) contain a positive fraction of forests or grasslands without land-use or land cover change during 1992–2020. To fulfill (1), predicted pixels include those whose predictor values lie within the input data range without outliers. The pixels that are excluded from the study domain because of each out-of-range predictor are shown in different colors. For each land cover type, all colored 0.5° pixels (both within the study domain or and excluded because of extrapolation) fulfill (2) by having a positive areal proportion that have not experienced land-use land cover change during 1992–2020 (Method). The white regions are areas not applicable to the land cover or excluded because of land-use land cover change during 1992–2020.

**Fig. S2.**
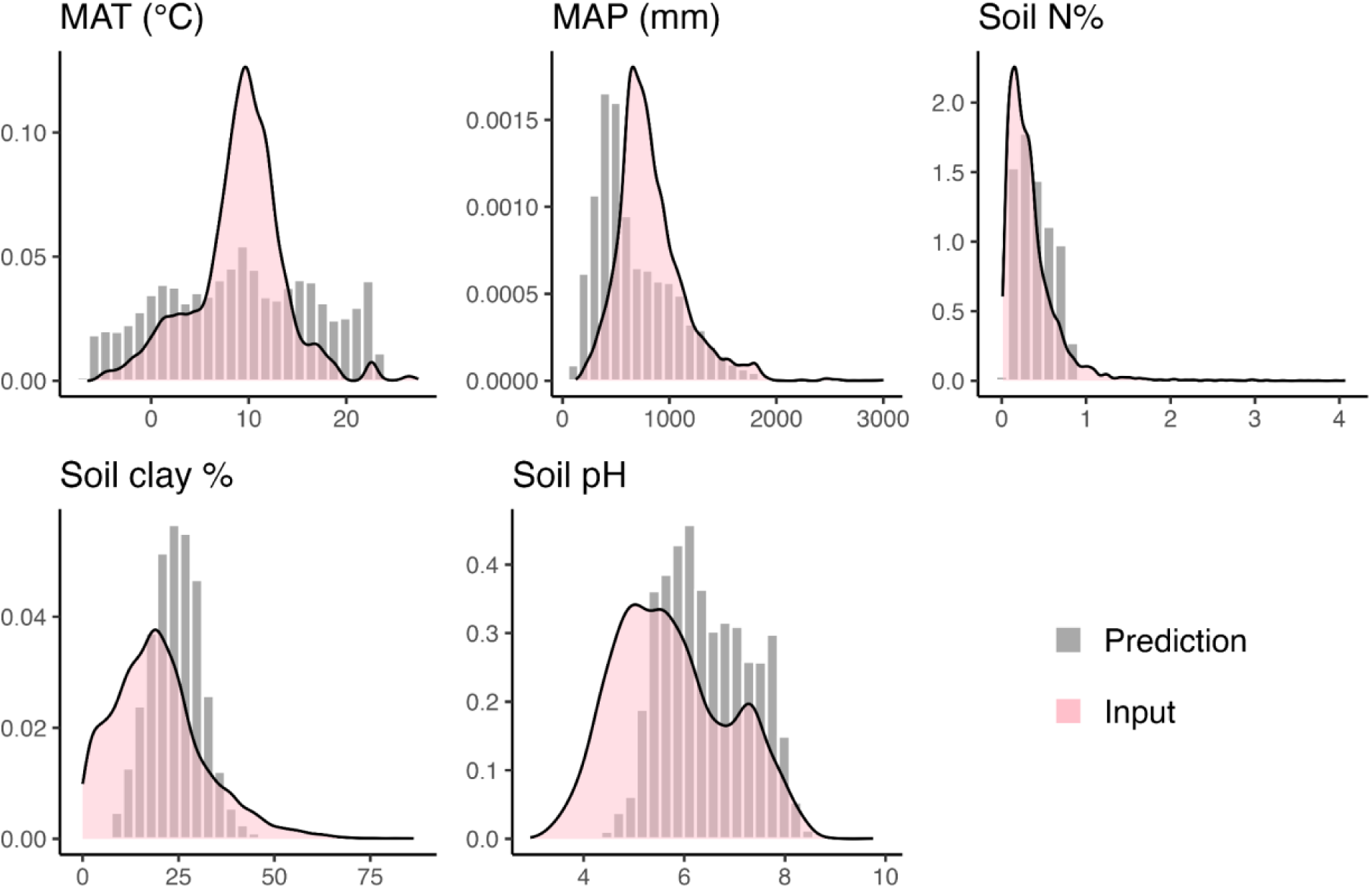
Density distribution of predictor values of predicted pixels and input data. The distribution of input data predictor values is well spread across those of the predicted pixels. Mean annual temperature (MAT) and mean annual precipitation (MAP) of the predicted years were used in the actual analysis. Here, we present the values of mean annual temperature (MAT) and mean precipitation (MAP) predictor layers in 1992 as examples.

**Fig. S3.**
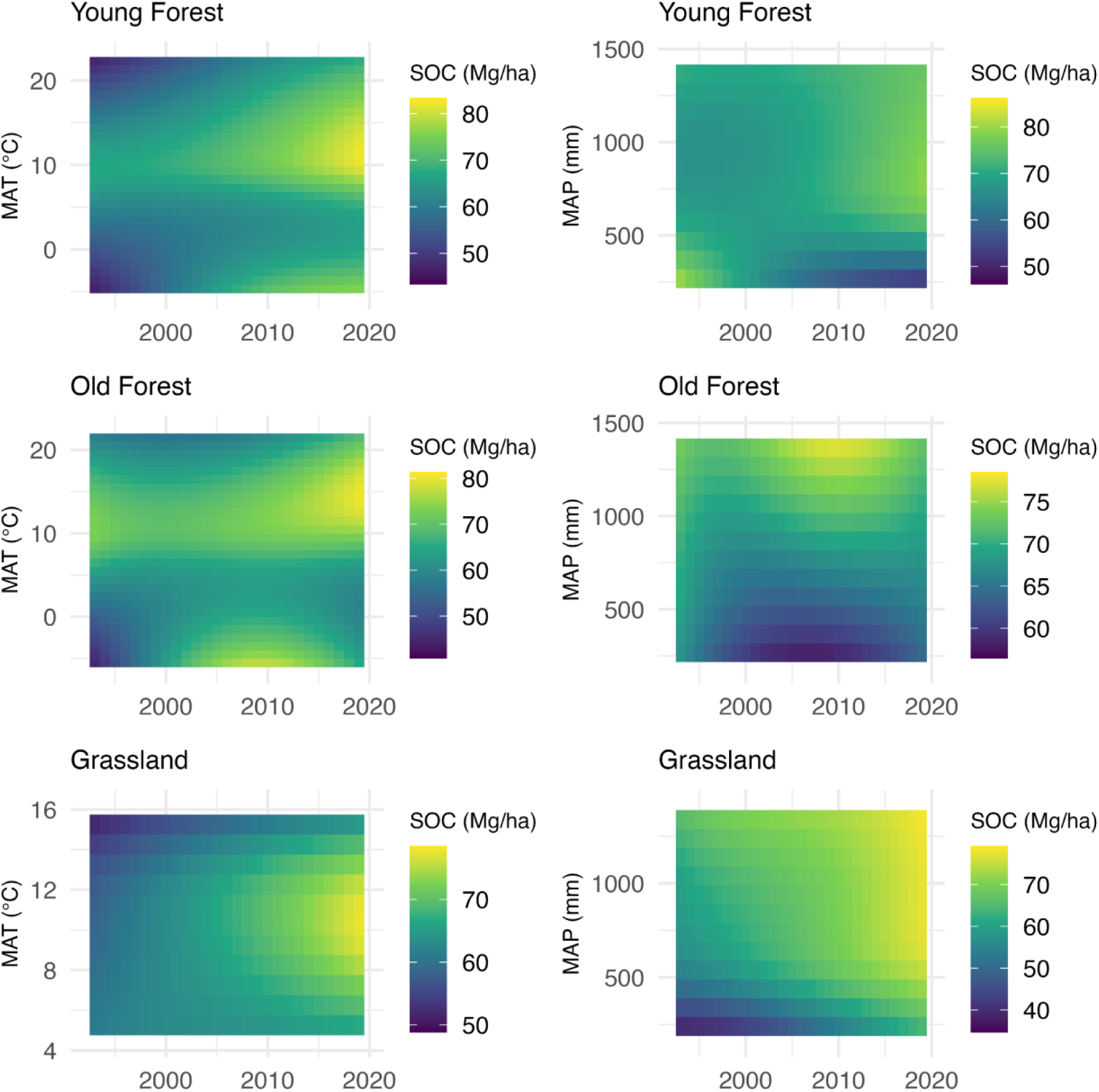
Heatmaps showing relationships between MAT, MAP, and year fitted in the statistical model (GAMM). In each panel, the heatmap color represents the predicted SOC stock (Mg C ha^-1^) under different MAT or MAP over the years. The SOC stock on these heatmaps predicted by the GAMM using average predictors values from the input data, while varying MAT or MAP. For each land cover, the MAT and MAP ranges plotted are the ranges covered in the study domain.

**Fig. S4.**
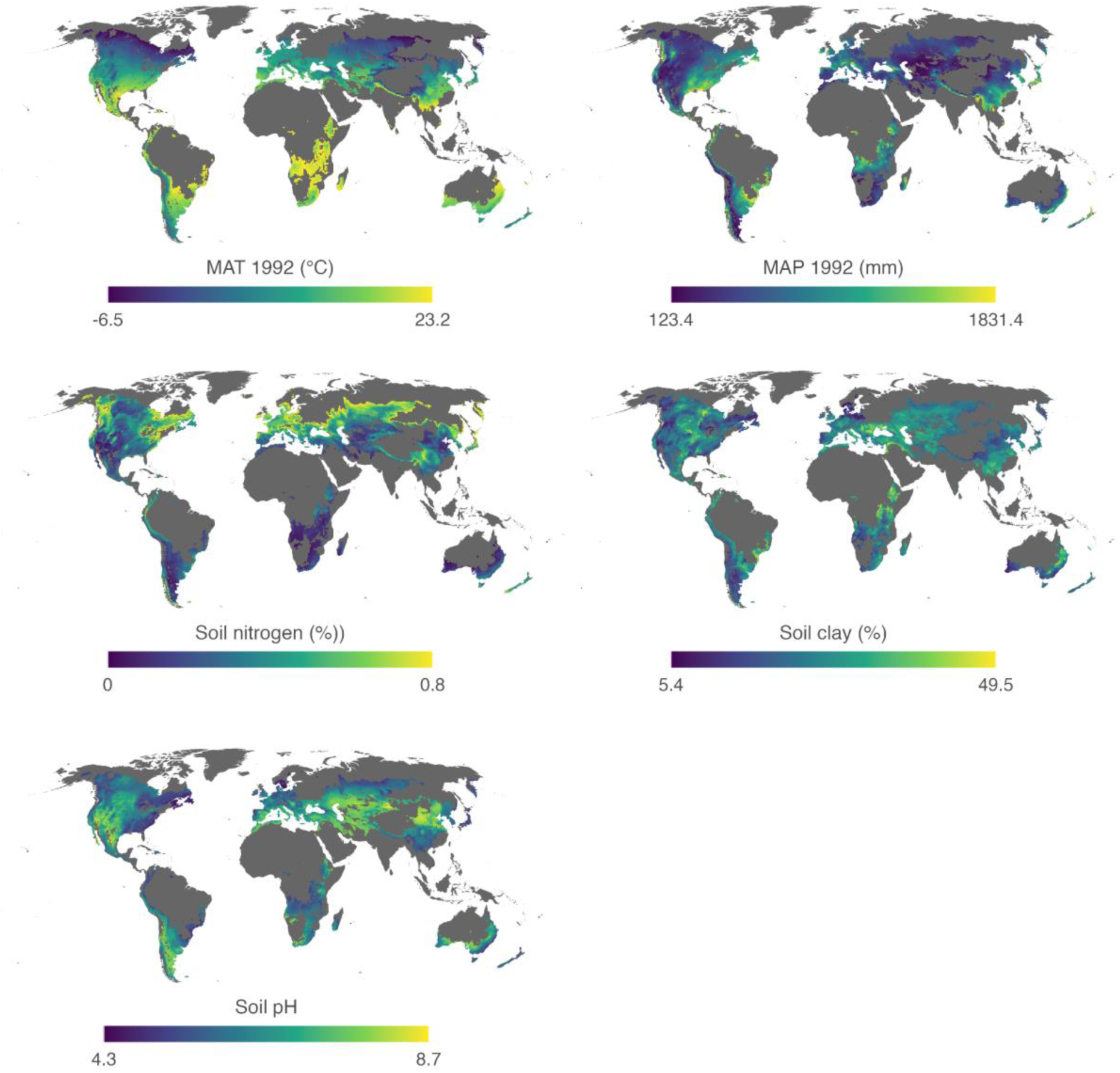
Values of predictors in all in-domain pixels. Mean annual temperature (MAT) and mean annual precipitation (MAP) of the predicted years were used in the actual analysis. Here we present MAT and MAP in 1992 as examples.

**Fig. S5.**
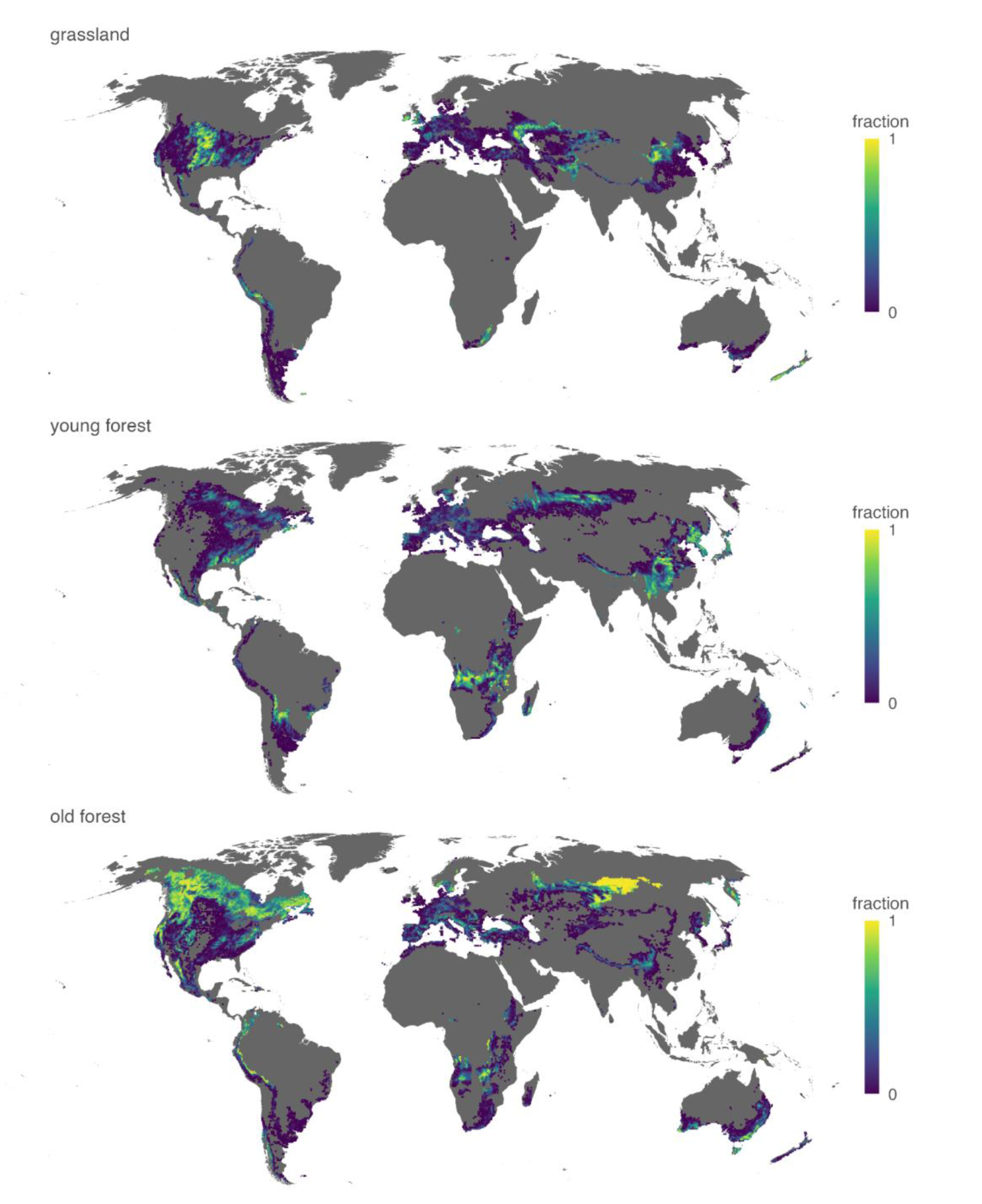
The area fractions of each land cover type in in-domain pixels. Calculations for land cover fractions are described in Methods.

**Table S1.**
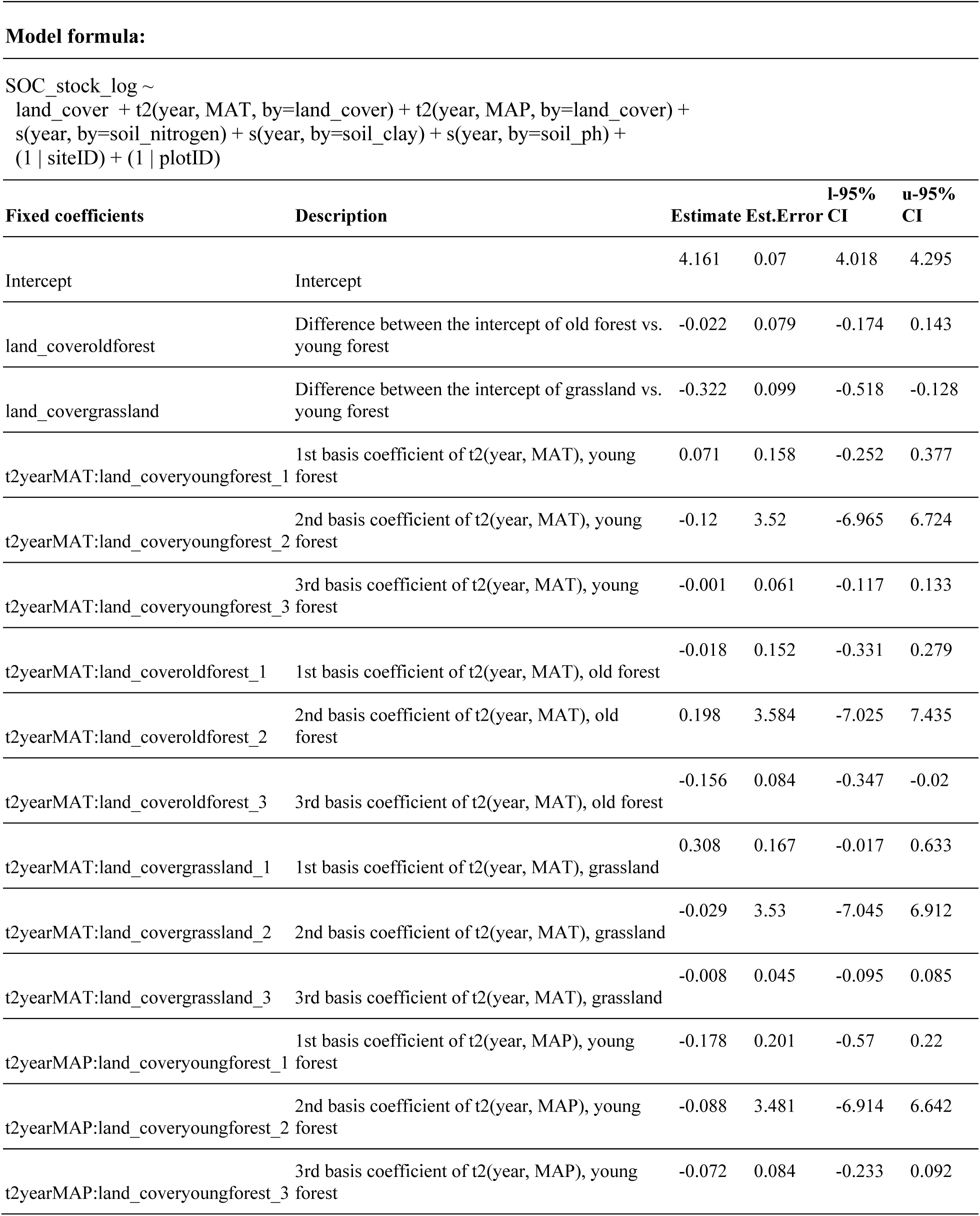
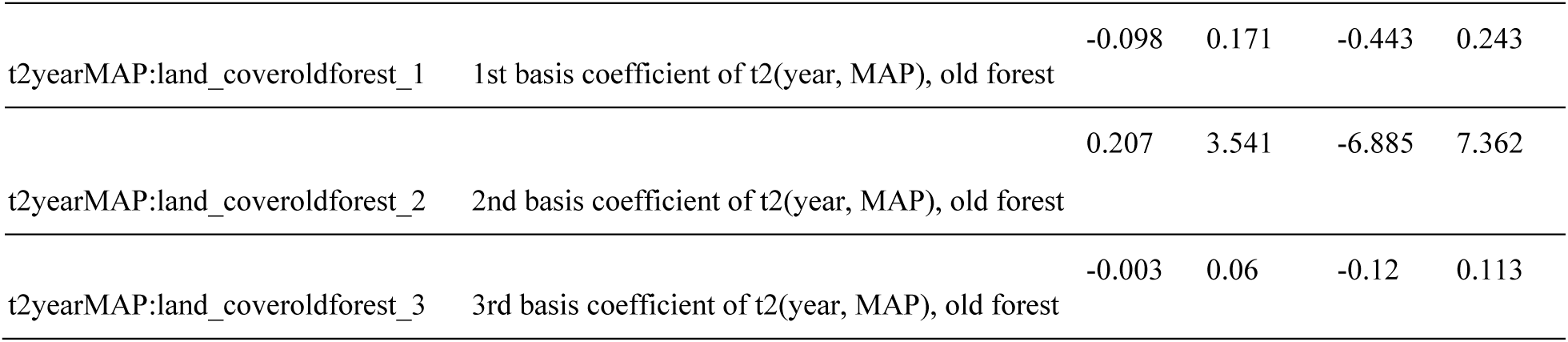
Summary output of fixed coefficients of the GAMM model. Est.Error: estimated standard error. I-95% CI: lower bound of 95% credible interval. u-95% CI: upper bound of 95% credible interval.

**Table S2.** TRENDY model estimates of SOC change in old forests and grasslands during 1992–2020 over the study’s spatial domain. SOC is estimated to be a sink if the change is positive, or a source if the change is negative.

| **Rank (from low to high)** | **Model** | **SOC change during 1992–2020 (Pg C yr<sup>-1</sup>)** |
|---|---|---|
| 1 | LPJwsl | -0.0402 |
| 2 | LPJ-GUESS | 0.014 |
| 3 | LPJmL | 0.0216 |
| 4 | SDGVM | 0.0235 |
| 5 | JULES | 0.0317 |
| 6 | ISAM | 0.0454 |
| 7 | E3SM | 0.0543 |
| 8 | DLEM | 0.0601 |
| 9 | LPX-Bern | 0.0651 |
| 10 | JSBACH | 0.0706 |
| 11 | CLASSIC | 0.0708 |
| 12 | ORCHIDEE | 0.0792 |
| 13 | OCN | 0.0998 |
| 14 | ISBA-CTRIP | 0.106 |
| 15 | IBIS | 0.1313 |
| 16 | VISIT | 0.1562 |
| 17 | EDv3 | 0.1833 |

**Table S3.**
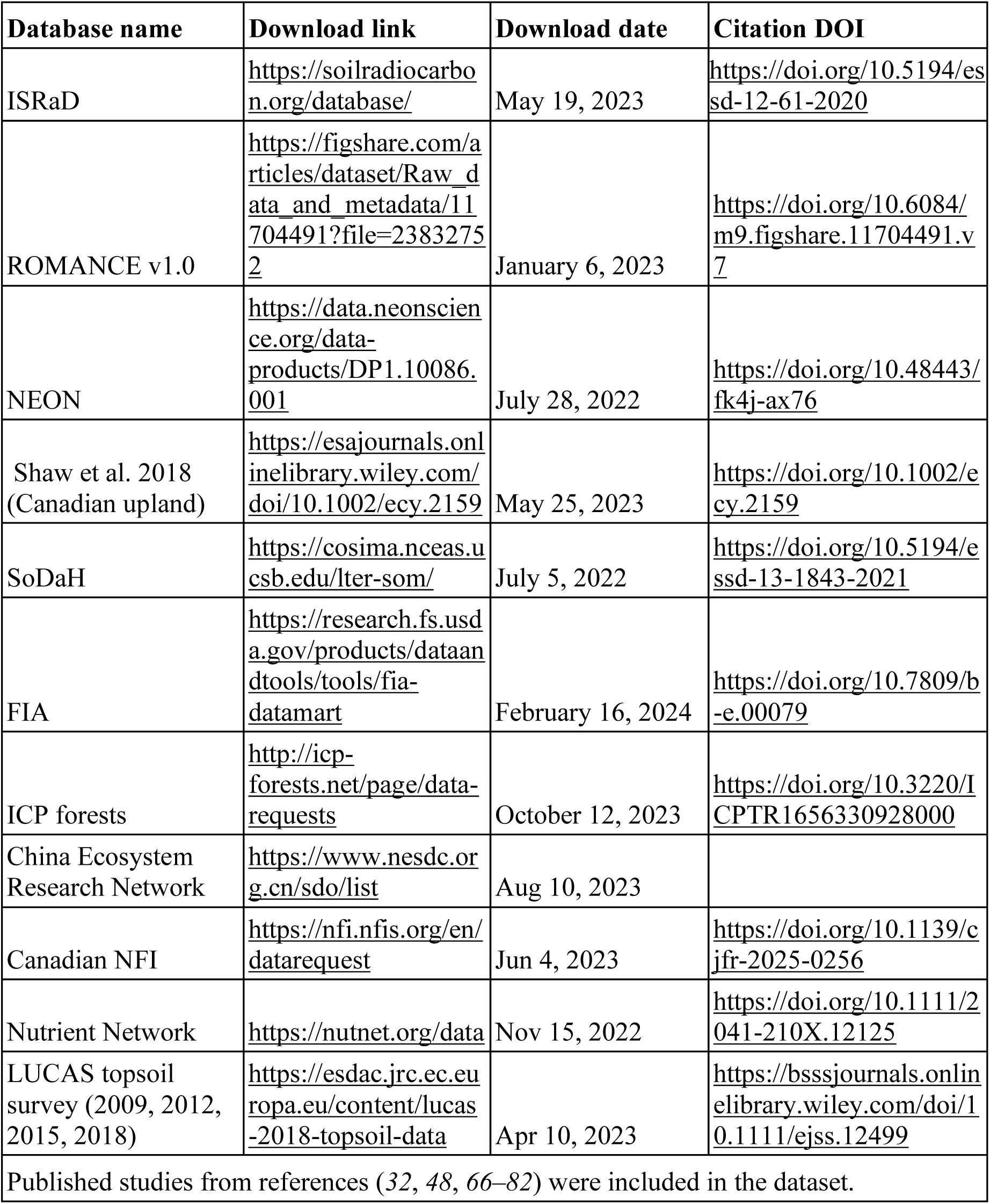
Data source details.

| Database name | Download link | Download date | Citation DOI |
|---|---|---|---|
| ISRaD | https://soilradiocarbon.org/database/ | May 19, 2023 | https://doi.org/10.5194/essd-12-61-2020 |
| ROMANCE v1.0 | https://figshare.com/articles/dataset/Raw_data_and_metadata/11704491?file=23832752 | January 6, 2023 | https://doi.org/10.6084/m9.figshare.11704491.v7 |
| NEON | https://data.neonscience.org/data-products/DP1.10086.001 | July 28, 2022 | https://doi.org/10.48443/fk4j-ax76 |
| Shaw et al. 2018 (Canadian upland) | https://esajournals.onlinelibrary.wiley.com/doi/10.1002/ecy.2159 | May 25, 2023 | https://doi.org/10.1002/ecy.2159 |
| SoDaH | https://cosima.nceas.ucsb.edu/lter-som/ | July 5, 2022 | https://doi.org/10.5194/essd-13-1843-2021 |
| FIA | https://research.fs.usda.gov/products/dataandtools/tools/fia-datamart | February 16, 2024 | https://doi.org/10.7809/b-e.00079 |
| ICP forests | http://icp-forests.net/page/data-requests | October 12, 2023 | https://doi.org/10.3220/ICPTR1656330928000 |
| China Ecosystem Research Network | https://www.nesdc.org.cn/sdo/list | Aug 10, 2023 |  |
| Canadian NFI | https://nfi.nfis.org/en/datarequest | Jun 4, 2023 | https://doi.org/10.1139/cjfr-2025-0256 |
| Nutrient Network | https://nutnet.org/data | Nov 15, 2022 | https://doi.org/10.1111/2041-210X.12125 |
| LUCAS topsoil survey (2009, 2012, 2015, 2018) | https://esdac.jrc.ec.europa.eu/content/lucas-2018-topsoil-data | Apr 10, 2023 | https://bsssjournals.onlinelibrary.wiley.com/doi/10.1111/ejss.12499 |
| Published studies from references (*32*, *48*, *66–82*) were included in the dataset. |  |  |  |

